# A forward genetic approach to mapping a *P*-element second site mutation identifies *DCP2* as a novel tumor suppressor in *Drosophila melanogaster*

**DOI:** 10.1101/2020.03.30.016865

**Authors:** Rohit Kunar, Rakesh Mishra, Lolitika Mandal, Debasmita P. Alone, Shanti Chandrasekharan, Jagat Kumar Roy

## Abstract

The use of transposons to create mutants has been the cornerstone of *Drosophila* genetics in the past few decades. Transpositions often create second-site mutations, devoid of transposon insertion and thereby affect subsequent phenotype analyses. In a *P*-element mutagenesis screen, a second site mutant was discovered on chromosome 3 wherein the homozygous mutant individuals show the classic hallmarks of mutations in tumor suppressor genes including brain tumour and lethality, hence the mutant line was initially named as *lethal (3) tumorous brain* [*l(3)tb*]. Classical genetic approaches relying on meiotic recombination and subsequent complementation with chromosomal deletions and gene mutations mapped the mutation to CG6169, the mRNA decapping protein 2 (*DCP2*), on the left arm of the third chromosome (3L), and thus the mutation was renamed as *DCP2*^*l(3)tb*^. Fine mapping of the mutation further identified the presence of a *Gypsy*-LTR like sequence in the 5’UTR coding region of *DCP2*, alongwith expansion of the adjacent upstream intergenic AT-rich sequence. The mutant phenotypes are rescued by Introduction of a functional copy of *DCP2* in the mutant background, thereby establishing the causal role of the mutation and providing a genetic validation of the allelism. With the increasing repertoire of genes being associated with tumor biology this is the first instance that the mRNA decapping protein is being implicated in *Drosophila* tumourigenesis. Our findings therefore imply a plausible role for mRNA degradation pathway in tumorigenesis and identify *DCP2* as a potential candidate for future explorations of cell cycle regulatory mechanisms.

## Introduction

With increasing interest in studies of classical tumor suppressors (Papagiannouli and Mechler, 2013; Ivanov et al., 2010), the search for new candidate proteins in tumor suppression has garnered importance (Tipping and Perrimon, 2013). In *Drosophila, P*-element mutagenesis provides a convenient method to identify, isolate and clone tagged genes while probing for genes which could be mutated to tumor formation (Mechler, 1994). Although identification and subsequent molecular analysis is convenient with *P*-element transpositions, second–site mutations devoid of any *P*-element insertion may be created during transposition (Liebl et al., 2006). In a *P*-element mutagenesis screen, a second site mutant was discovered wherein the homozygous mutant individuals showed prolonged larval life, developed larval brain tumors with increased number of superficial neuroblasts and abnormal chromosomal condensation along with overgrowth in the wing and the eye-antennal discs and were lethal in the larval/pupal stages. Since all these are hallmarks of mutations in tumor suppressor genes in *Drosophila* (Gateff and Schneiderman, 1969; Gateff E, 1974; Gateff E, 1978), the mutation was named as *l(3)tb* [*l(3)tb: lethal (3) tumorous brain*] owing to its location on the third chromosome and the phenotypes manifested. Genetic and molecular analyses mapped the mutation to *DCP2* on the left arm of chromosome 3 (cytogenetic position 72A1) and hence the allele was named as *DCP2*^*l(3)tb*^. While complementation analyses of the mutation with alleles of *DCP2* show phenotypes similar to *l(3)tb* homozygotes and confirm the proposed allelism, over-expression of wild type *DCP2* in the mutant background rescues the mutant phenotypes, thereby providing a genetic validation of allelism. Subsequent fine mapping identified the presence of a *Gypsy*-LTR like sequence in the 5’UTR coding region, downstream to the transcription start site (TSS) of *DCP2. DCP2* codes for the mRNA decapping protein 2, which belongs to the NUDIX family of pyrophosphatases and was identified almost a decade ago through a yeast genetic screen (Dunckley and Parker, 1999). Being one of the major components of the decapping complex, DCP2 is conserved in worms, flies, plants, mice, and humans (Wang et al., 2002). DCP2 is activated by DCP1 and they function together as a holoenzyme to cleave the 5’ cap structure of mRNA (LaGrandeur and Parker, 1998; Coller and Parker, 2004; Parker and Song, 2004; She et al., 2008). *DCP2*^*l(3)tb*^ bears an incomplete LTR sequence from the *gypsy* element and develops brain tumors in *Drosophila*, thereby demanding considerable exploration of the exact perturbations in the DNA–protein interactions caused by its presence. Although mRNA decapping plays a significant role in mRNA turnover and translation, widely affecting gene expression (Mitchell and Tollervey, 2001; Raghavan and Bohjanen, 2004; Song et al., 2010), simultaneous links between mRNA degradation genes, retrotransposons and tumors have not been observed and/or investigated so far. Therefore, the novel allele *DCP2*^*l(3)tb*^ reveals a new perception for functional roles of mutant lesions and the ensuing perturbations in gene regulation in tumor biology.

## Materials and Methods

### Fly strains and rearing conditions

All flies were raised on standard agar-cornmeal medium at 24±1°C. *Oregon R*^+^ was used as the wild type control. The *l(3)tb* mutation (*yw;* +*/*+; *l(3)tb* /*TM*6*B, Tb*^*1*^, *Hu, e*^*1*^) was isolated in a genetic screen and the mutation was maintained with the *TM6B* balancer. The multiply marked “*rucuca*’’ (*ru h th st cu sr e ca/TM6B,Tb*) and “*ruPrica*’’ (*ru h th st cu sr e Pr ca/TM6B,Tb*) chromosomes were employed for recombination mapping (Lindsley and Zimm 1992). *w; Δ2-3, Sb/TM6B, Tb*^*1*^, *Hu, e*^*1*^ (Cooley *et al.* 1988) and *CyO, P{Tub-Pbac/T}2/Wg*^*Sp-1*^;+*/TM6B, Tb, Hu, e*^*1*^ were used for providing transposase source for *P* element and *piggyBac* specific transposable element, respectively, in mutagenesis experiment. The *y*^*1*^*w; P{Act5C-GAL4}25F01/CyO* and *yw;* +*/*+; *Tub-GAL4/ TM3, Sb, e*, were obtained from the Bloomington *Drosophila* Stock Center. The lethal insertion mutants of gene *DCP2, viz., PBac{RB}DCP2*^*e00034*^*/TM6B, Tb*^*1*^ *Hu, e*^*1*^ (Thibault *et al.* 2004) and *P{GT1}DCP2*^*BG01766*^*/TM3, Sb*^*1*^, *e*^*1*^ (Lukacsovich *et al.* 2001) were obtained from Exelixis Stock Center, Harvard University and Bloomington *Drosophila* stock center, respectively.

Deficiency stock *Df(3L)RM96* was generated in the laboratory (for details of characterisation, refer to **Supplementary Table S3**) using progenitor *P* element stocks viz. *P{RS5}5-SZ-3486, P{RS5}5-SZ-3070, P{RS3}UM-8356-3, P{RS3}UM-8241-3, P{RS3}CB-0072-3, yw P{70FLP, ry*^+^*}3F*^*iso*^*/y*^+^*Y; 2*^*iso*^; *TM2/TM6C, Sb, w*^*1118*^ _*iso*_*/y*^+^*Y; 2*^*iso*^; *TM2/TM6C, Sb* obtained from Vienna *Drosophila* Resource Center (Golic and Golic, 1996; Ryder *et al.* 2007). Various deficiency stocks and transposon insertion fly stocks (**Supplementary Tables S1 and S2**) used for complementation analysis were obtained from Bloomington *Drosophila* stock centre and Exelixis stock centre.

### Analysis of lethal phase in *l(3)tb* homozygotes

For analysis of lethal phase and morphological anomalies associated with the homozygous *l(3)tb* mutation, embryos were collected at the intervals of 2h on food filled Petri dishes. Embryos from wild type flies were collected as controls. The total number of eggs in each plate was counted and the embryos were allowed to grow at 23°C or 18°C or 16°C (±1°C). Hatching of embryos and further development of larval stages was monitored to determine any developmental delay. Mutant larvae, at different stages, were dissected and the morphology of larval structures was examined.

### Identification of Mutant Locus in *l(3)tb*

#### a. Meiotic recombination mapping of *l(3)tb* mutation

Genetic recombination with multiple recessive chromosome markers, *ru cu ca*, was performed to map mutation in *y w:* +*/*+; *l(3)tb/TM6B, Tb* mutant. The *y w; l(3)tb/TM6B* males were crossed to virgin +*/*+; *ru Pri ca/TM6B* females to recover *l(3)tb* without *y w* on X-chromosome. The F1 *l(3)tb/TM6B* males were crossed to virgin +*/*+; *ru cu ca* females and the F2 progeny +*/*+; *l(3)tb/ru cu ca* virgin females were selected. These F2 virgins were then crossed to *ru Pri ca/TM6B* males to score the frequency of recombinants in the F3 progeny. Thereafter, all the F3 progeny males obtained, were individually scored for *ru, h, th, st, cu, sr, e* and *ca* phenotypes and then they were individually crossed with virgin *l(3)tb/TM6B* females to identify which of them had the *l(3)tb* mutation along with other scored markers.

#### b. Complementation mapping of the *l(3)tb* mutation

Complementation analysis of the mutation in *l(3)tb* allele was carried out in two stages. Firstly, deficiency stocks spanning the entire chromosome 3 (**Supplementary Table S1**) were used to identify the mutant loci, and secondly, lethal *P*-insertion alleles selected from the region narrowed down through recombination and deficiency mapping (**Supplementary Table S2**) were harnessed to further identify the mutant gene(s) in *l(3)tb*. In either case, virgin females of *yw;* +*/*+; *l(3)tb/TM6B,Tb* were crossed with the males of the various deficiency stocks and/or the lethal *P*-insertion alleles and the non-tubby F1males heterozygous for *l(3)tb* and the deficiency were scored for the phenotype(s).

Reversion analysis was performed by the excision of *piggyBac* transposon in *DCP2*^*e00034*^ with the help of *piggyBac* specific transposase source, *CyO, P{Tub-Pbac}2/Wg*^*SP-1*^ (Thibault *et al.* 2004) or by the excision of *P*-element in *DCP2*^*BG01766*^ strain using transposase from the ‘jumpstarter’, Δ*2-3,Sb/TM6B, Tb*^*1*^, *Hu, e*^*1*^. Virgin flies from the ‘mutator stocks’, *viz., DCP2*^*e00034*^ or *DCP2*^*BG01766*^ strain were crossed to male flies from respective ‘jumpstarter stock’. F1 male flies with mosaic eye pigmentation carrying both the transposase and respective transposons were selected and crossed to JSK-3 (*TM3, Sb, e*^*1*^*/TM6B, Tb*^*1*^, *Hu, e*^*1*^) virgins and from the next generation (F2), rare white eyed revertant flies were selected (Figure S1).

### Fine Mapping of *l(3)tb* mutation

#### a. Genomic DNA Isolation, PCR and Southern hybridisation

Genomic DNA for polymerase chain reaction (PCR) was isolated by homogenizing 50 male flies from each of the desired genotype or 80-100 third instar larvae from homozygous mutant *l(3)tb* (Sambrook *et al.* 1989). Based on the results obtained from genetic mapping, identification of the candidate region in *DCP2* was done by overlapping PCR based screening, wherein the entire genomic span of *DCP2* was amplified using 28 primer pairs from 3L:15811834..15819523 (**Supplementary Tables S4, S5, S6**) (Rozen and Skaletsky 2000). After identifying the candidate region, it was validated with the primer pair, Dbo_F: 5’-ACAACATTCACTCCATGGAACACCT-3’ and DCP2_P19_R: 5’-TGCTCACCGAACTTTTTCGCGATCT-3’. The primer pair DCP2_F: 5’-ATAACAAAAAAGTTATGGTACCACCCCCGCGTTGTATTCT-3’ and DCP2_R: 5’-AGATTTCGATGTATATGGATCCGTCCCAACCTTTGCGTCT-3’ was designed to amplify the full length gene along with flanking sequences (500 bp on either side). In either case, the thermal cycling parameters included an initial denaturation at 96°C (2 min) followed by 30 cycles of 30 s at 94°C, 45 s at 72°C, and 15 min at 68°C. Final extension was carried out at 68°C for 20 min. The PCR products were electrophoresed on 0.8% agarose gel with O’GeneRuler 1kb plus DNA ladder (Thermo Scientific, USA). An 812 bp region (3L: 15825979..15826790) spanning the candidate mutated region in *DCP2* was PCR amplified and ligated in pGEM-T vector (Promega) to generate the pGEM-T-812 clone. The ∼430 bp fragment isolated during primer walking (see below) was purified and ligated in pTopo-TA-XL vector (Invitrogen, USA) to generate the pTopo-TA-XL-430 clone. Digestion, ligation and transformation were performed using standard protocols as described in Sambrook and Russell, 2001. Southern hybridizations were performed according to Sambrook and Russell, 2001. Following electrophoresis and gel pre-treatments, DNA was transferred on to positively charged nylon membranes (Roche, Germany). Hybridizations were performed at 68°C with 0.02% SDS, 5X SSC, 0.5% Blocking reagent, and 0.1% laurylsarcosine with probes generated from pGEM-T-812 and pTopo-TA-XL-430 plasmids. DIG Labelling and chemiluminescent detection were performed as per the manufacturer’s instructions (Roche, Germany).

#### b. Sequencing, Primer Walking and CNV detection

To confirm the fidelity of amplification automated DNA Sequencing was performed (ABI – 3130, USA) as per the manufacturer’s instructions. Primer walking was initiated with the primers Dbo_F and DCP2_P19_R and from the terminal part of the sequence obtained, new primers P19_W2_F 5’-GGAGATCTGTTTGAAATATCTCTTCACATT–3’ and P19_W2_R 5’– GGCGCGTCAGCATTGTTCATACAAAGCTAC-3’ were designed. Long-range PCR with P19_W2 was performed as described previously. Sequence chromatograms were assessed and analyzed with FinchTV 1.4.0, Geospiza Inc. Semi-quantitative assessment of copy number variance (CNV) of the intergenic sequence in *DCP2*^*l(3)tb*^ was determined through PCR analyses. A 156 bp sequence (3L: 15826497..15826652) was chosen to be amplified by CNV_F 5’-ACAGTTGGCTCTGTGATAAATGT-3’ and CNV_R 5’-AGTGCAACGGAAGGGAATCT-3’ against an internal control sequence of 153 bp, corresponding to the gene *Dsor*, amplified by the primer pair. Thermal cycling parameters included an initial denaturation at 95°C (5 min) followed by 28 cycles of 30 s at 94°C, 30 s at 60°C, and 30 s at 72°C. Final extension was carried out at 72°C for 10 min. The PCR products were electrophoresed on 2% agarose gel with a 100-bp DNA ladder (BR Biosciences, India).

### Immunocytochemistry

The imaginal discs and/or brain ganglia were collected from wild type *Oregon R*^+^ wandering 3^rd^ instar larvae, just before pupation (110 h, AEL) and in mutant homozygous *l(3)tb* from day 6 and day 10/12. The tissues were processed for immunostaining as described in Banerjee and Roy, 2018, with the desired antibodies. Primary antibodies used in this study were - Anti-Discs large, 4F3 (1:50, Developmental Studies Hybridoma Bank, Iowa,USA), Anti-Armadillo (1:100, a kind gift by Prof LS Shashidhara, IISER Pune, India), Anti-Elav (Rat-Elav-7E8A10, 1:100, DSHB, USA), Anti-DE-Cadherin (DCAD2, 1:20, DSHB, Iowa, USA), Anti-phospho-Histone 3 (1:500, Millipore, Upstate, USA), Anti-Deadpan (1:800, a kind gift from Prof. Volker Hartenstein, University of California, USA) and Anti-Cyclin E (HE12; sc-247, 1:50, Santa Cruz, India). Appropriate secondary antibodies conjugated either with Cy3 (1:200, Sigma-Aldrich, India) or Alexa Fluor 488 (1:200; Molecular Probes, USA) or Alexa Fluor 546 (1:200; Molecular Probes, USA) were used to detect the given primary antibody, while chromatin was visualized with DAPI (4’, 6-diamidino-2-phenylindole dihydrochloride, 1μg/ml Sigma-Aldrich). Counterstaining was performed with either DAPI (4’, 6-diamidino-2-phenylindole dihydrochloride, Sigma) at 1µg/ml, or phalloidin-TRITC (Sigma-Aldrich, India) at 1:200 dilutions. Tissues were mounted in DABCO (antifade agent, Sigma). The immunostained slides were observed under Zeiss LSM 510 Meta Laser Scanning Confocal microscope, analysed with LSM softwares and assembled using Adobe Photoshop 7.0.

### Statistical analysis

Sigma Plot (version 11.0) software was used for statistical analyses. All percentage data were subjected to arcsine square-root transformation. For comparison between the control and experimental samples, One-Way ANOVA was performed. Data were expressed as mean ± S.E. of mean (SEM) of several replicates.

## Results

### *l(3)tb* homozygotes show the classic hallmarks of cancer in *Drosophila* including developmental delay, abnormal karyotype, larval/pupal lethality alongwith tumorous brain and wing imaginal disc

Developmental analysis of *l(3)tb* homozygotes showed that while embryos hatched normally and developed alike their heterozygous siblings [*l(3)tb*/*TM6B*], the third instar larvae reached the wandering stage quite late with the larval stage extending up to 12 or 13 days (**Figure 1B**). Although 66.8% of the larvae survived to pupate (**Table 1**), they died in the pupal stage following bloating, enhancement in size and cessation of growth (**Figure 1A**). Hence, the mutation is absolutely lethal with the lethality being pronounced in the pupal stage. Lowering the temperature to 16°C or 18°C reduced the larval mortality, causing 96% of larvae to pupate but did not improve pupal survival (**Figure 1 C and D**). Analysis of larval brain and imaginal discs in the homozygotes in the early (Day 6) and late (Day 10-12) larval phase showed gross morphological alterations in the size of the larval brain, wing and eye imaginal discs (**Figure 2A–G**) as compared to the wandering wild type third instar larvae (115h ALH; **<u>A</u>**fter **<u>L</u>**arval **<u>H</u>**atching). The brain was smaller in size than the wild type (*Oregon R*^+^) or heterozygous [*l(3)tb*/*TM6B*] individuals till 115 ALH but started showing aberrant growth in the dorsal lobes thereafter, showing significant differences in the diameter and area of the lobes. The overgrown brain hemispheres remained more or less symmetric in most of the cases, except in some where it got deformed and fused with the imaginal discs (**Figure 2 J and K**). A similar trend in morphological aberration was observed in the wing discs, which remained smaller initially but enlarged sufficiently later (**Figure 2L**), with abnormal protrusion in the wing pouch. Analysis of mitotically active cell population by screening for the metaphase marker protein, phosphorylated histone H3 (PH3) revealed increased number of active mitoses in the mutant homozygous brains (**Figure 3A-O; 3V**) and wing discs (**Figure 3P-V**) (Day 6) in comparison to the wild type, the number of which increased with increase in larval age of the mutant. However, mitotic karyotypes of the mutant brain lacked numerical aberrations, despite showing extensive variability in condensation (**Figure 2 H and I**).

**Table 1.** Homozygous mutation in *l(3)tb* causes larval and pupal lethality.

| Genetic cross<br>(23<sup>0</sup>C<sub>±</sub>1) | Total<br>Eggs | No. of eggs<br>hatched | 3<sup>rd</sup> Instar larvae<br>transferred | Pupae<br>formed | Flies<br>eclosed |
| --- | --- | --- | --- | --- | --- |
| ***l(3)tb/TM6B***<br>**X**<br>***l(3)tb/TM6B*** | 1500 | 972<br>(O= 64.8%)<br>(E=66.7%) | Non-Tubby-468<br>(O=48.1%)<br>(E= 50%) | Non-tubby<br>313<br>(O=66.8%) | Non Tubby<br>0 |
|  |  |  | Tubby<br>473<br>(O= 48.6%)<br>(E=50%) | Tubby<br>468<br>(O=98.9%)<br>(E=100%) | Tubby<br>465<br>(O=99.4%)<br>(E=100%) |

Numbers in parenthesis indicate the percentage observed (O) and expected (E) values out of the total progeny from previous stage.

**Figure 1.**
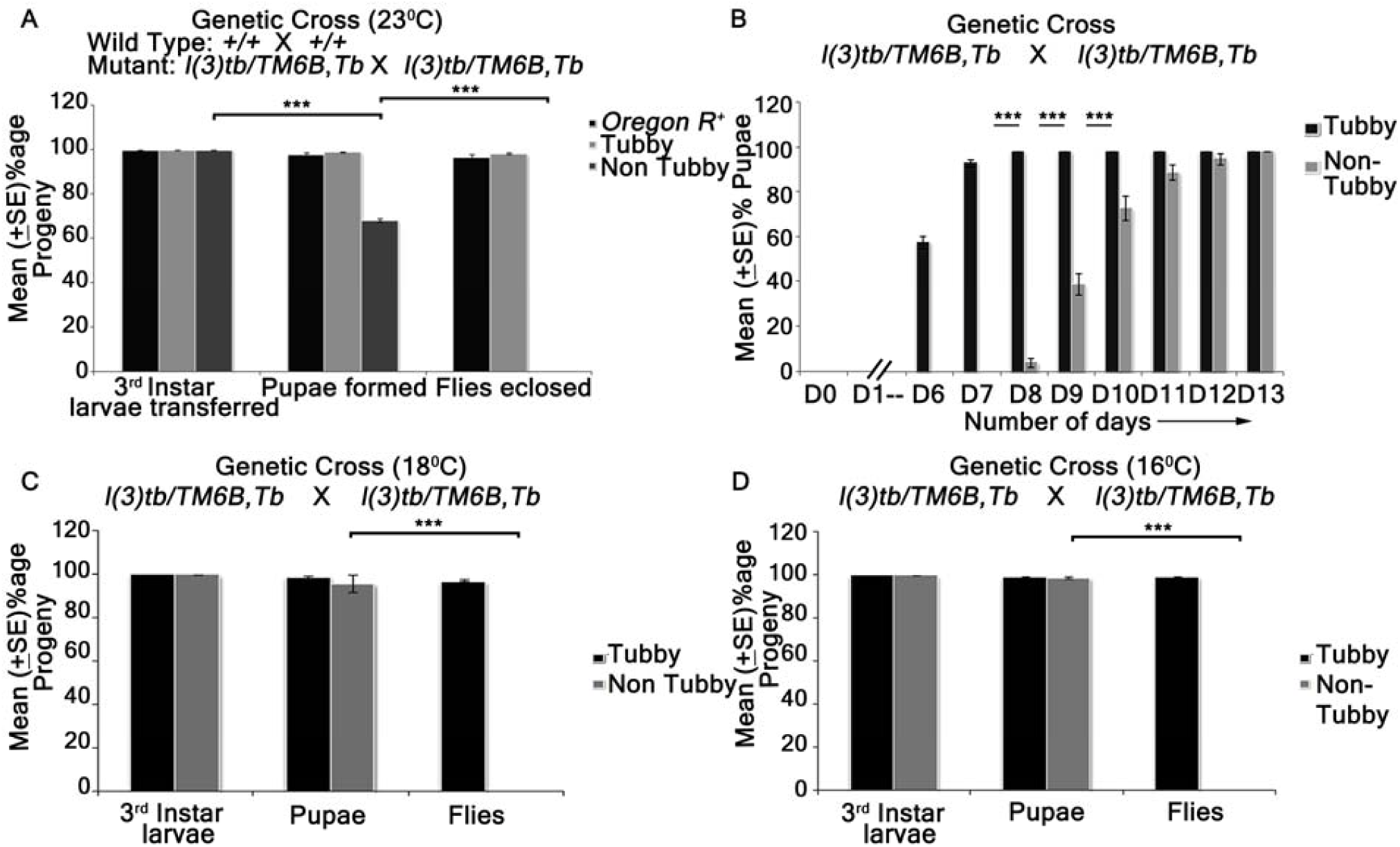
Homozygous *l(3)tb* show delayed larval development with lethality at larval/pupal stage (A, B) and is not a conditional temperature sensitive allele (A, B, C). Homozygous *l(3)tb* progeny, at 23°C, showed lethality at larval and pupal stages and no flies eclosed as compared to wild type and heterozygous *l(3)tb* progeny with balancer chromosome (A). Homozygous *l(3)tb* progeny individuals demonstrated extended larval life up to day 12/13 where as heterozygous progeny individuals followed the normal wild type pattern of development (B). (C) and (D) show significant increase in viability of homozygous (non-tubby) *l(3)tb* larvae at lowered temperatures of 18°C and 16°C respectively, though there also occurred absolute lethality at pupal stages. Each bar represents mean (±S.E.) of three replicates of 100 larvae in each. *** indicates p<0.005 *** indicates p<0.005.

**Figure 2.**
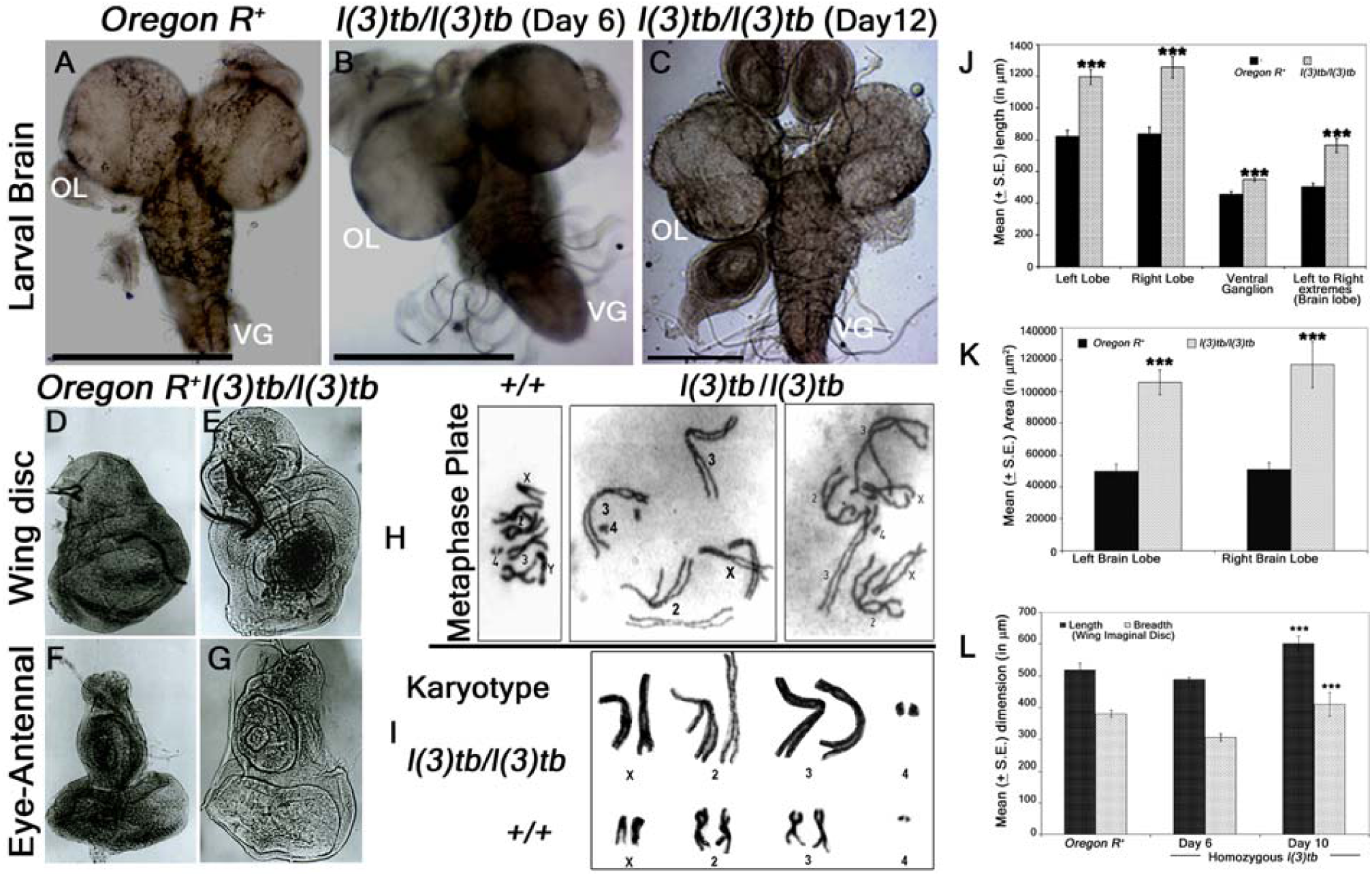
Homozygous *l(3)tb* mutants show severe morphological alteration in delayed 3^rd^ instar larval brain, wing and eye-antennal disc of. Homozygous *l(3)tb* mutant 3^rd^ instar larvae revealed tumorous brain of day 12 (C) as compared to day 6 of homozygous mutant (B) and day 5 of wild type, *Oregon R*^+^ (A). *l(3)tb* homozygotes exhibited highly significant differences in the overall circumference of the left and right brain lobes in the delayed stage (day 10) as compared to the respective wild type brain lobes (J). Significant differences were found in the area (µm^2^) of respective brain lobes of *l(3)tb* homozygotes and wild type (K). Dimension of wing and eye-antennal imaginal discs of delayed 3^rd^ instar larvae from homozygous *l(3)tb* mutant revealed significant increase in size (D,E,F,G). Length and breadth of wing discs from 3^rd^ instar larvae of *l(3)tb* mutant of day 6, was found to be smaller than the wing imaginal discs from wild type, but wing discs from extended larval period (day 10) showed significant increase in the size (L). Metaphase chromosome preparation of brain cells (H) from wild type and *l(3)tb* homozygotes exhibited abnormal karyotypes (I) where *l(3)tb* homozygotes showed less condensed and extended chromosome morphology as compared to wild type, *Oregon R*^+^. *** denotes p<0.005

**Figure 3.**
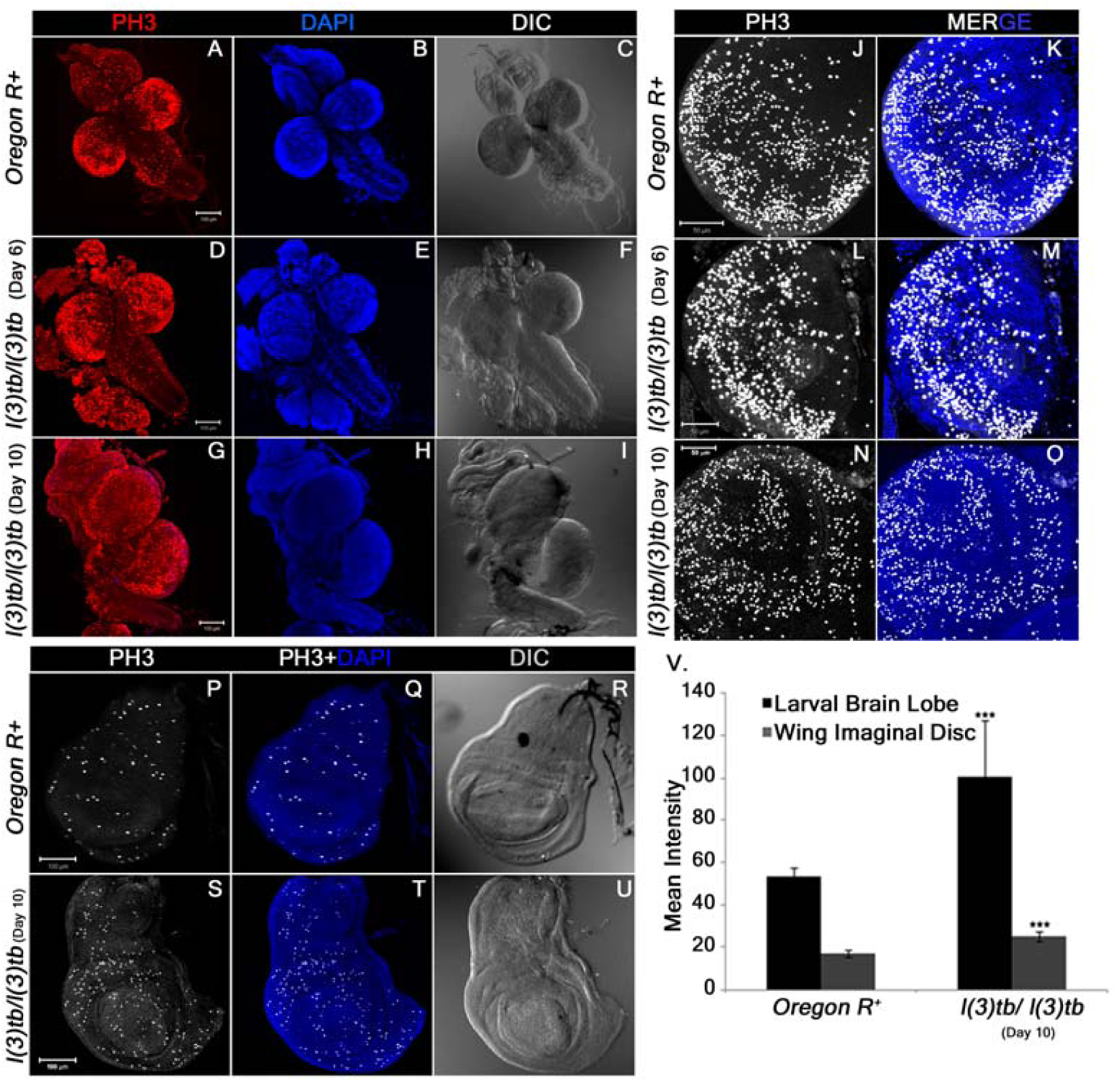
Enhanced mitotic potential observed in the tumourous tissues of homozygous *l(3)tb* as shown in larval whole brain (A), brain lobes (D, G) and wing imaginal discs (S) immunostained with phosphor-histone 3 (PH3), a potent mitotic marker. Distribution of PH3 labeled cells counter stained with DAPI cells in wild type (A) and homozygous *l(3)tb* (Day 6 and Day 10) larval brain (D, G) and also in wild type brain lobes (B, C) and homozygous mutant brain lobes (E, F for day 6; H, I for day 10) indicated high mitotic index as compared to wild type. Similarly, more mitotic positive cells were seen in tumorous wing imaginal discs (day 10) of homozygous mutant *l(3)tb* (S) as compared to wild type, *Oregon R*^+^ (P). DIC images (C, F, I and R, U) illustrates external normal morphology in wild type and more pronounced tumorous phenotypes in homozygous *l(3)tb* larval brain and wing imaginal discs. Quantitative analysis showed increase in the number of mitotic positive cells in homozygous mutant larval brain lobes and wing imaginal discs as compared to wild type and difference was highly significant (V). The images are projections of optical sections acquired by confocal microscopy. Staining was done in triplicates with 10 brains and 15 wing imaginal discs in each group. Significant difference is represented as *** *P*≤0.005 using one-way ANOVA.

### Eye-antennal discs and leg imaginal discs also show morphological and developmental anomalies in *l(3)tb* homozygous individuals

Global analysis of morphological aberrations in the mutant homozygotes showed that besides the tumorous brain and wing imaginal discs, eye-antennal discs and leg imaginal discs were also overgrown with a transparent appearance. Expression of Elav and Dlg in the eye-antennal discs revealed similarities to the developmental perturbations observed in the wing discs and brain. In the early third instar mutant larvae (Day 6), all photoreceptor cells showed expression of Elav, similar to the wild type tissue (**Figure 4J and N**). However, during advanced stages of larval tumorigenesis (Day 10), it dwindled eventually (**Figure 4R**). The Elav expressing cells which are posterior to the morphogenetic furrow co-express Dlg and demonstrate the typical ommatidial arrangement. In the mature mutant larvae however, the eye discs demonstrate significant deviations from the normal regular arrangement of ommatidia. The leg imaginal discs, which reside in close proximity to the brain and wing imaginal discs also show enlargement in size which increases with advancement and retention of larval stage. They show gradual disruption of normal expression of DE-cadherin and Armadillo (**Figure 5**), alike tumorous wing discs (see above), implying the mutation and subsequent tumor to affect developmental homoeostasis in adjacent tissues as well.

**Figure 4.**
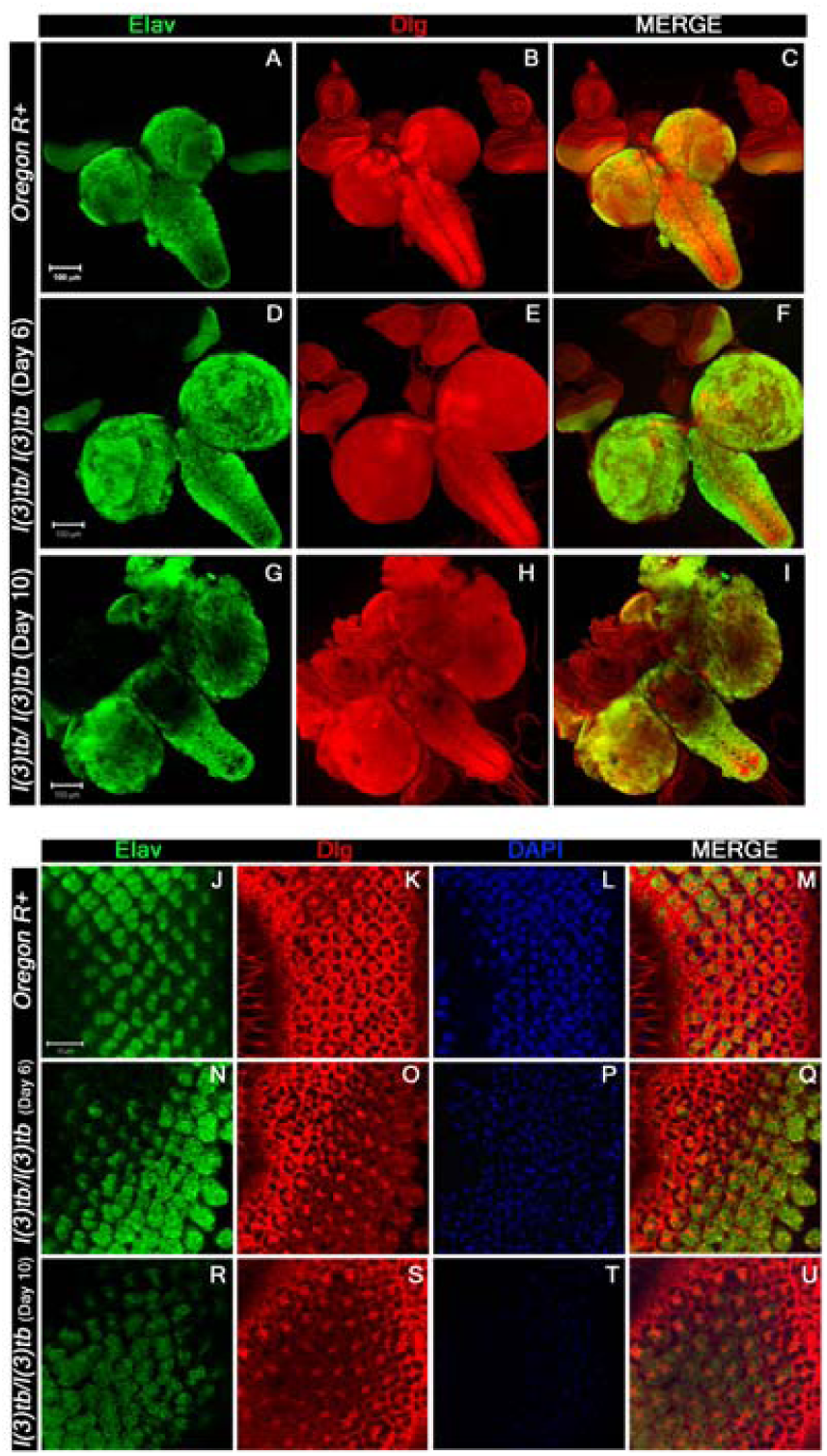
Confocal photomicrograph show loss of mature neurons and increase in junctional protein, Dlg, in delayed (Day 10) homozygous *l(3)tb*. 3^rd^ instar larval brain shows intense staining of Elav (green) in day 6 (D) of homozygous mutant later on show loss of staining in enlarged brain of day 10 (G), while the wild type brain (A) showed normal pattern of Elav staining. Dlg stained the ventral nerve chord and central brain in optic lobes of wild type (B), which is similar in day 6 of homozygous mutant brain (E) but in delayed larval brain, day 10, the pattern was altered (H). Scale shown is 100µm. Neuronal tissue from eye imaginal discs also display loss of neurons seen through Elav staining in day 10 (R) as compared to day 6 (N) in homozygous *l(3)tb* mutant as well as to wild type (J). Pattern of junctional protein, Dlg, in eye imaginal discs is also altered in day 10 (S) as compared to day 6 (O) and wild type (K). Counter stain with DAPI shows very weak intensity in day 10 (T) reflecting disintegrating chromatin as compared to day 6 (P) and wild type (L). Scale bar represents 10µm.

**Figure 5.**
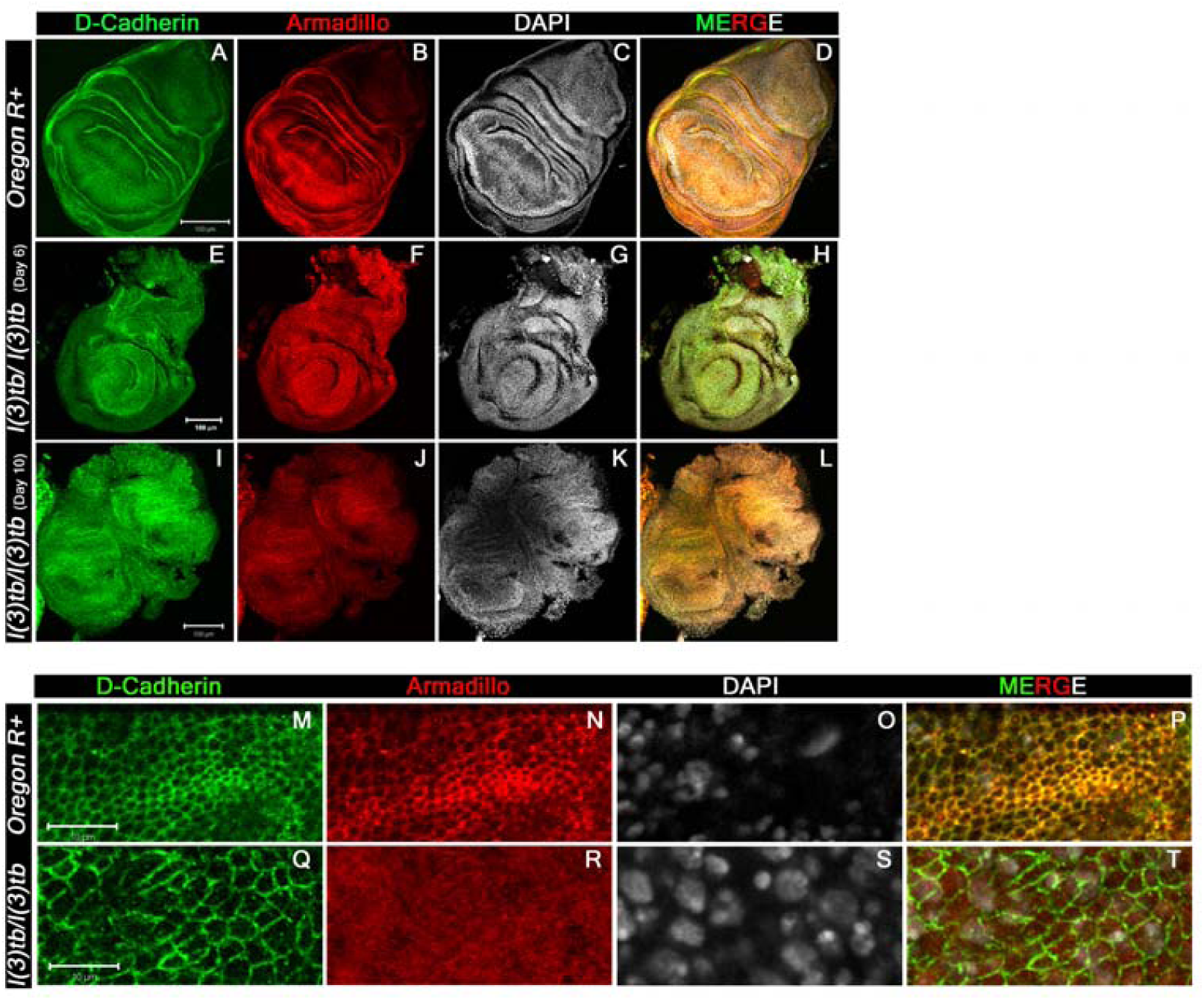
Confocal images of 3^rd^ instar larval wing imaginal discs immunolabeled to visualize the altered distribution pattern of cadherin-catenin complex proteins. Tumor caused in the homozygous *l(3)tb* mutant completely altered the distribution pattern of both, trans-membranous protein DE-cadherin (A, E, I, M, Q) and Armadillo (β-Catenin, B, F, J, N, R) adheren junctional proteins. Alteration of both proteins is more pronounced in the wing imaginal discs from mutant larva during extended larval life (I, J) than in the early wing imaginal disc (E, F) as compared to distinct pattern of DE-cadherin (A) and Armadillo (B) in the wild type wing imaginal discs. Armadillo is a binding partner of trans-membranous protein DE-cadherin having roles in cell adhesion and regulate tissue organization and morphogenesis. Merged images also substantiate the altered distribution of both junctional proteins in the homozygous mutant (H, L) as compared to the wild type (D) where co-localization is indicated by yellow pattern. Higher magnification of wing imaginal disc (pouch region) demonstrate altered distribution pattern of DE-cadherin (Q) and Armadillo (R) in homozygous *l(3)tb* mutant as compared to wild type (N, R). Increase in cell size seen in homozygous *l(3)tb* mutant (Q) as compared to wild type (M). Complete loss of Arm staining observed in homozygous *l(3)tb*)(R) whereas normal pattern seen in wild type wing disc (N). Chromatin size also altered in homozygous *l(3)tb* (S) as compared to wild type (O). Wild type shows clear co-localization of D-Cad and Arm (P), while there is complete loss of co-localization in homozygous *l(3)tb* wing imaginal discs (T). Scale bar represents 100 µm (A to L) and 10 µm (M to T).

### Genetic mapping through meiotic recombination and complementation mapping identify *l(3)tb* to be allelic to *DCP2*

The mutation *l(3)tb*, being recessive and on the third chromosome, was maintained with *TM6B* balancer. Analysis of meiotic recombination frequencies of an unmapped mutation with known markers is a classical technique that has been routinely employed to identify its cytogenetic position. In order to bring *l(3)tb* in a chromosome with such markers (*ru cu ca*), we allowed meiotic recombination to occur between *l(3)tb* and the 8 recessive markers present on the “*rucuca*” chromosome (**Table 2**). 113 recombinant males were observed and recombination frequencies were calculated in centiMorgan (cM). **Table 3** shows the recombination frequencies of each marker (locus) with the mutation *l(3)tb*. Preliminary analysis suggested that *l(3)tb* was close to *thread* (*th*) with minimum recombination events between the two loci (2.65%). Further analysis of recombination events between *h-l(3)tb* [17.78%], *st-l(3)tb* [1.23%] and *cu-l(3)tb* [8.29%] (**Table 4**) and comparing with the positions of each of the markers, the mutation was estimated to be located left of *thread* (43.2 cM; band 72D1) between 41.71 cM–42.77 cM, *i.e.*, in the cytological position 71F4-F5.

**Table 2.**
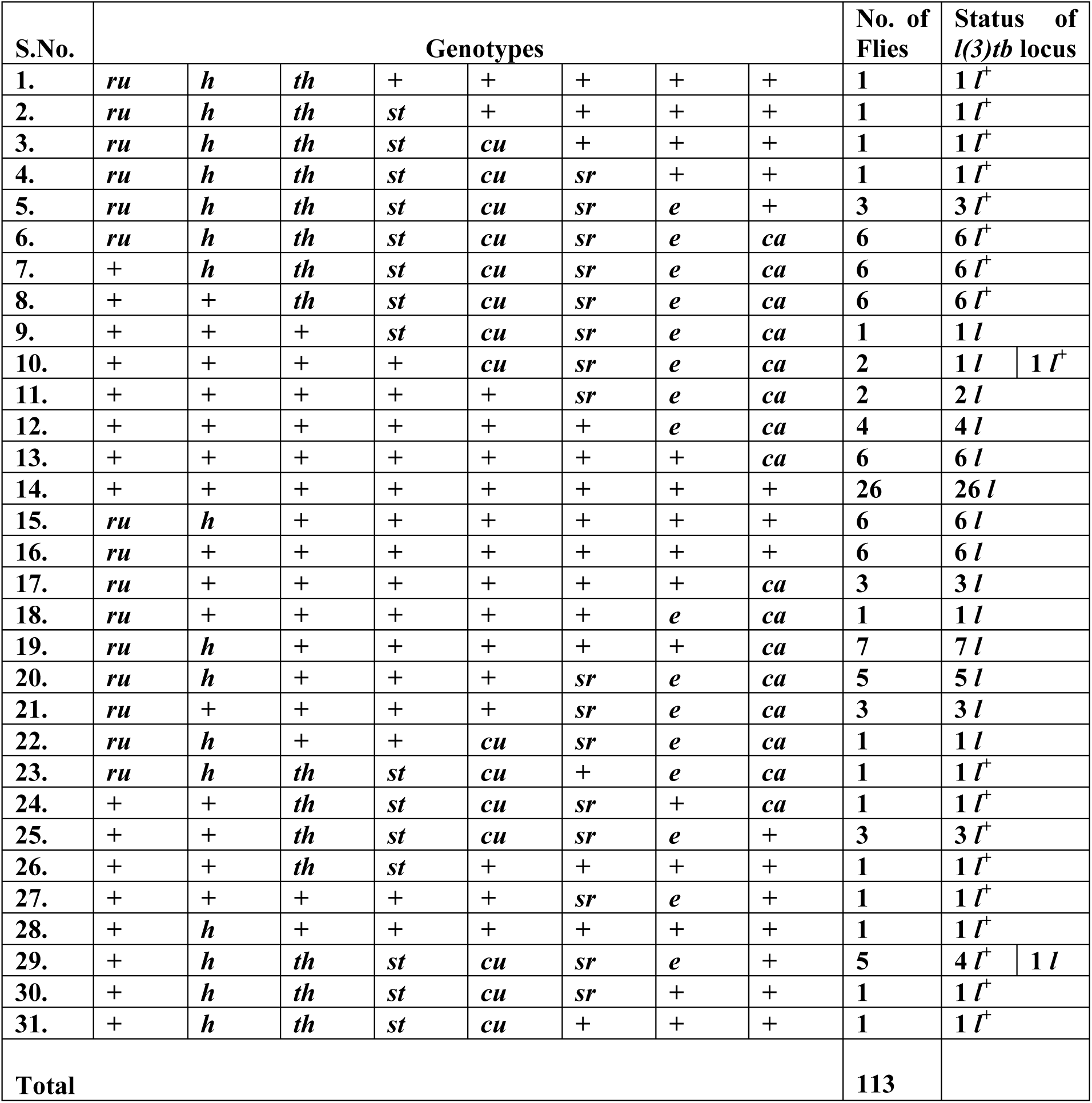
Rearranged genotypes of 113 males after various recombination events between all the eight visible markers of *rucuca* chromosome.

**Table 3.** Recombination frequencies (RF) between various recessive markers on *rucuca* chromosomes (*roughoid, hairy, thread, scarlet, curled, stripe, ebony*, and *claret*) and *l(3)tb*.

| Sl. No. | Association of marker with *l(3)tb* | Flies | | | | Recombination frequency (RF)<br>$\frac{R}{P+R} \times 100$ |
| --- | --- | --- | --- | --- | --- | --- |
|  |  | Parental (P) |  | Recombinant (R) |  |  |
|  |  | Genotype | No. of flies | Genotype | No. of flies |  |
| 1. | *ru - l* | *ru*<sup>+</sup> *l*<br>*ru* *l*<sup>+</sup> | 18<br>} 62<br>44 | *ru*<sup>+</sup> *l*<sup>+</sup><br>*ru* *l* | 18<br>} 51<br>33 | 45.13 |
| 2. | *h - l* | *h*<sup>+</sup> *l*<br>*h* *l*<sup>+</sup> | 54<br>} 79<br>25 | *h*<sup>+</sup> *l*<sup>+</sup><br>*h* *l* | 12<br>} 34<br>22 | 30.08 |
| 3. | *th - l* | *th*<sup>+</sup> *l*<br>*th* *l*<sup>+</sup> | 74<br>} 110<br>36 | *th*<sup>+</sup> *l*<sup>+</sup><br>*th* *l* | 1<br>} 3<br>2 | 2.65 |
| 4. | *st - l* | *st*<sup>+</sup> *l*<br>*st* *l*<sup>+</sup> | 73<br>} 108<br>35 | *st*<sup>+</sup> *l*<sup>+</sup><br>*st* *l* | 2<br>} 5<br>3 | 4.42 |
| 5. | *cu - l* | *cu*<sup>+</sup> *l*<br>*cu* *l*<sup>+</sup> | 71<br>} 105<br>34 | *cu*<sup>+</sup> *l*<sup>+</sup><br>*cu* *l* | 3<br>} 8<br>5 | 7.07 |
| 6. | *sr - l* | *sr*<sup>+</sup> *l*<br>*sr* *l*<sup>+</sup> | 62<br>} 94<br>32 | *sr*<sup>+</sup> *l*<sup>+</sup><br>*sr* *l* | 4<br>} 19<br>15 | 16.8 |
| 7. | *e - l* | *e*<sup>+</sup> *l*<br>*e* *l*<sup>+</sup> | 56<br>} 86<br>30 | *e*<sup>+</sup> *l*<sup>+</sup><br>*e* *l* | 7<br>} 27<br>20 | 23.89 |
| 8. | *ca - l* | *ca*<sup>+</sup> *l*<br>*ca* *l*<sup>+</sup> | 42<br>} 74<br>32 | *ca*<sup>+</sup> *l*<sup>+</sup><br>*ca* *l* | 16<br>} 49<br>33 | 43.3 |

**Table 4.**
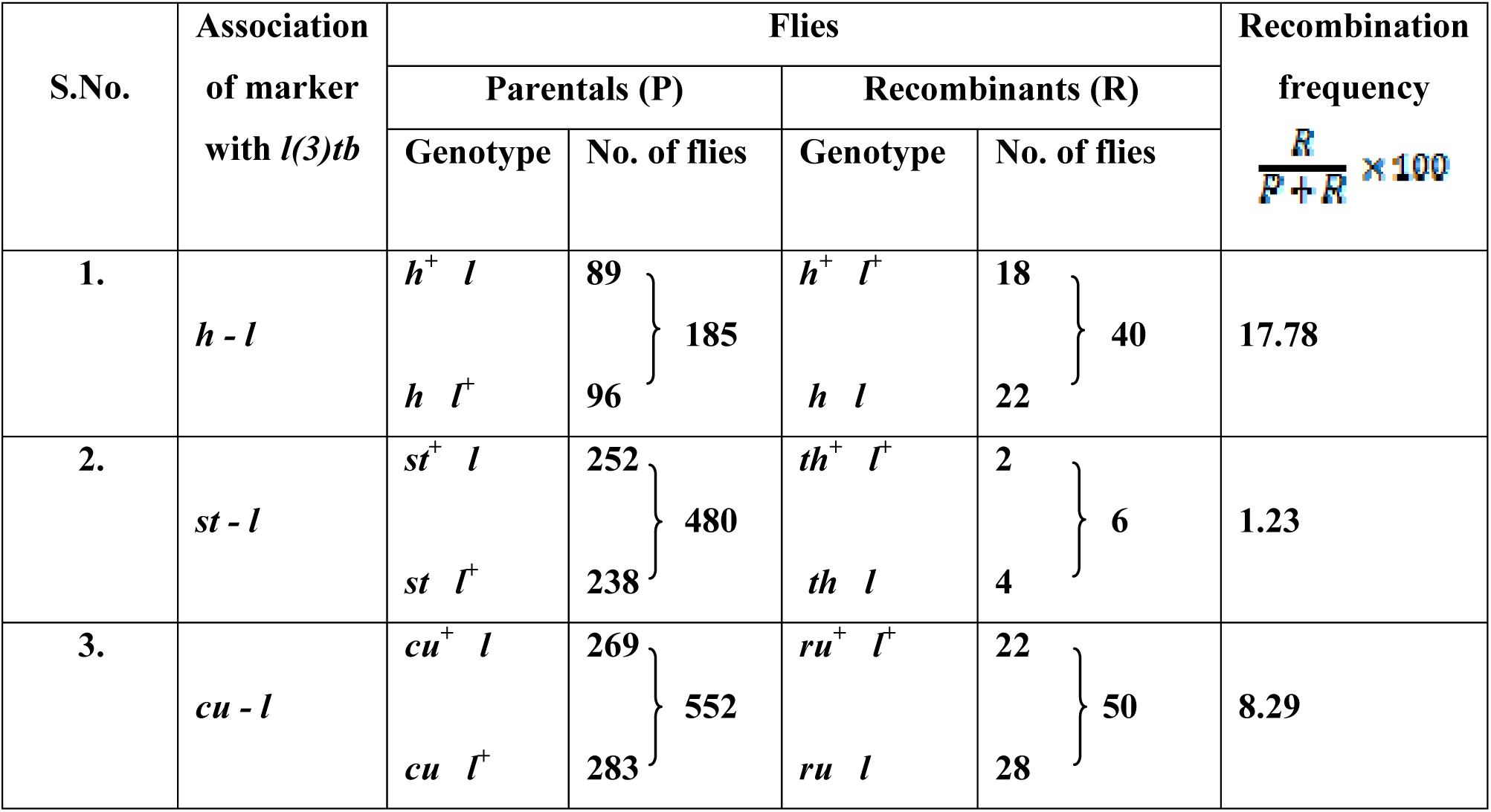
Recombination events between *h-l, st-l* and *cu-l*.

Complementation analysis with molecularly defined Drosdel and Exelixis deficiency lines (N=85), spanning the entire chromosome 3, identified four lines which failed to complement the mutation, *viz., Df(3L)BSC774, Df(3L)BSC575, Df(3L)BSC845* and *Df(3L)RM95*, which was generated in the lab using progenitor RS stocks. Trans-heterozygotes *l(3)tb*/*Df(3L)BSC575* were pupal lethal and the dying non-tubby larvae showed phenotypes similar to *l(3)tb* homozygotes, suggesting the mutation to reside between 71F1 and 72A1 on the left arm of chromosome 3. Further analysis using six deletion lines belonging to the above region (71F1–72A2) identified the mutation to reside between 71F4 to 71F5, which strangely is a gene desert region. Complementation analyses performed with lethal insertion alleles (N=26) of genes residing proximal or distal to 71F4-F5 identified two lethal *P*-element insertion alleles of *DCP2* (mRNA decapping protein 2; CG6169), *viz., P{GT1}DCP2*^*BG01766*^ and *PBac{RB}DCP2*^*e00034*^, which failed to complement the mutation *l(3)tb* (**Figure 7A and C**) as well as those deletions which had failed to complement *l(3)tb*, implying the mutation to be allelic to *DCP2* (72A1).

**Figure 6.**
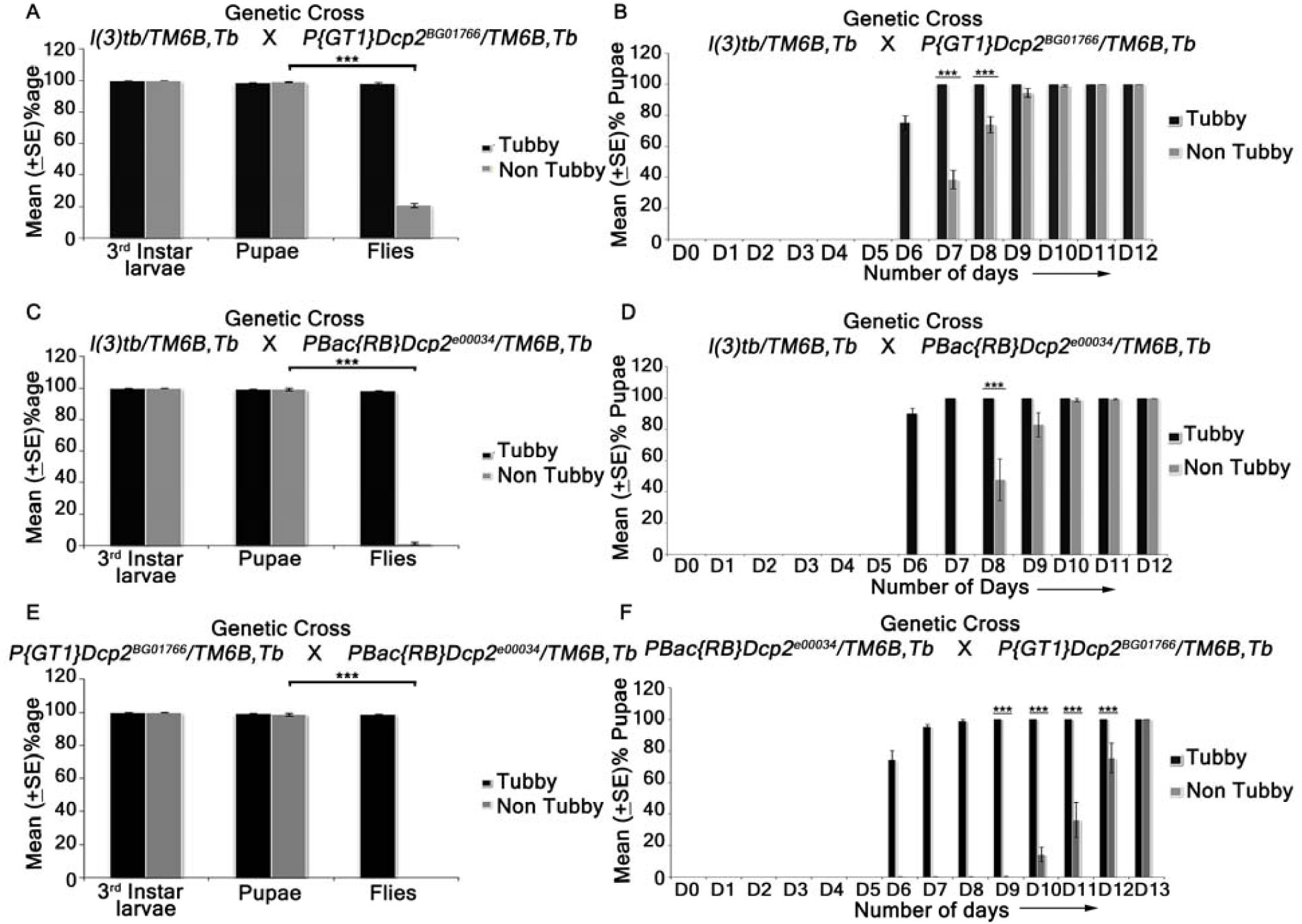
Viability assay performed on various hetero-allelic combinations between alleles of gene *DCP2* and the mutation in *l(3)tb*. Homozygous *l(3)tb* exhibited larval as well as pupal lethality. 69% of homozygous larvae pupated whereas no fly eclosed from the pupae. *l(3)tb* trans-heterozygous with *P{GT1}DCP2*^*BG01766*^ showed only 18.4% fly eclosed (A). *l(3)tb/ PBac{RB}DCP2*^*e00034*^ trans-heterozygote (C) causes 100% lethality at pupal stage. Trans-allelic combination *P{GT1}DCP2*^*BG01766*^/ */PBac{RB}DCP2*^*e00034*^ (E) also exhibited 100% pupal lethality. Developmental delay seen in trans-heterozygotes *l(3)tb /P{GT1}DCP2*^*BG01766*^ (B) and *l(3)tb/PBac{RB}DCP2*^*e00034*^ (D) as in homozygous *l(3)tb*. Progeny from heterozygous for both the alleles of *DCP2* gene, *PBac{RB}DCP2*^*e00034*^ */P{GT1}DCP2*^*BG01766*^ (F) also exhibited developmental delay. *** indicates p<0.005.

**Figure 7.**
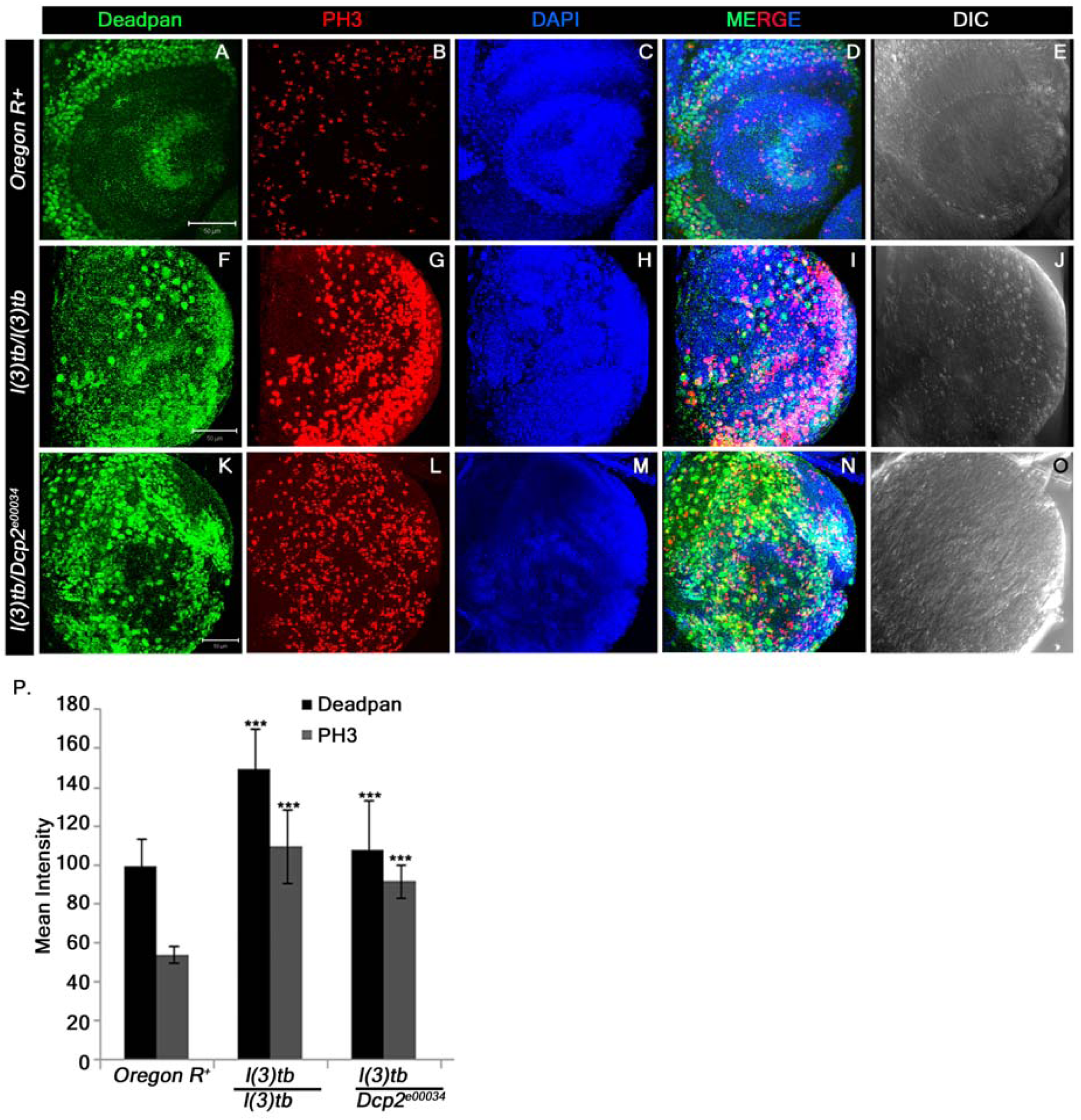
Heterozygous combination of *l(3)tb* with *DCP2*^*e00034*^ allele resulted in to significant increase in the number of neuroblasts and mitotically active cells. Confocal projection sections showing immunolocalisation of Deadpan, a neuroblast marker (Green, A, F, K) for picking neuroblasts and phosphohistone 3 (PH3, red, B, G, L) marking the mitotic cells are shown. Enhanced neuroblast population in homozygous mutant (F) and in heterozygous *l(3)tb* with *DCP2*^*e00034*^ allele (K) Similarly, increased number of mitotic cells (PH3 positive) also occurred in heterozygous *l(3)tb* with *DCP2*^*e00034*^ allele (L), similar to homozygous *l(3)tb* mutant (G). NBs and mitotic positive cells are quantified (P) and the differences are statistically significant when compared with wild type. *** *P* >0.005. Scale bar indicates 50µm.

### Trans-heterozygotes of *DCP2* mutants and *l(3)tb* show developmental delay, tumorous larval brain with elevated neuroblast numbers, larval/pupal lethality and developmental defects in escapee flies

Trans-heterozygotes of *l(3)tb* with either allele of *DCP2, viz., P{GT1}DCP2*^*BG01766*^ and *PBac{RB}DCP2*^*e00034*^, showed developmental delay. In either case, trans-heterozygous third instar larvae showed persistence of larval stage till Day 10 ALH (**Figure 6B and D**), and show tumorous phenotypes of brain and wing imaginal discs (**Supplementary Figure S4**), similar to the *l(3)tb* homozygotes. Expression pattern of Deadpan (Dpn), a marker for neuroblasts show increased number of neuroblasts in the larval brain of the trans-heterozygotes as well as *l(3)tb* homozygotes (**Figure 7F, K and P**). Also, the trans-heterozygous progeny showed a higher mitotic index as compared to the wild type progeny, similar to the *l(3)tb* homozygotes (**Figure 7G, L and P**). While *PBac{RB}DCP2*^*e00034*^*/l(3)tb* was found to be 100% pupal lethal, *P{GT1}DCP2*^*BG01766*^*/l(3)tb* was only 81.6% lethal (**Figure 6A and B**), with the rest 18.4% pupae eclosing as flies. However, the escapee flies showed several developmental abnormalities, *viz.*, defects in wing (9.5%), thorax closure (3.2%), loss of abdominal para-segments and abdominal bristles (3.2%), and presence of melanotic patches (22.2%), leg defects (41.3%) or eclosion defects (12.7%) (**Supplementary Figure S2**). Analysis of compound eyes in these escapees revealed complete loss of regular arrangement of ommatidia and ommatidial bristles (**Supplementary Figure S3**). Abnormal external genitalia were also observed in the male escapees (data not shown). Subsequent analysis of fertility showed that the trans-heterozygous escapee flies had compromised fertility with only 40% of the males and 21.7% of the females being fertile (**Table 5**).

**Table 5.** Fertility assay of trans-heterozygotes *P{GT1}DCP2*^*BG01766*^*/l(3)tb* demonstrating male and female sterility.

| **Cross** | $l(3)tb/P\{GT1\}DCP2^{BG01766}$<br>(males)<br>X<br>+/+<br>(Virgin females) | $l(3)tb/P\{GT1\}DCP2^{BG01766}$<br>(Virgin females)<br>X<br>+/+<br>(males) |
| --- | --- | --- |
| **Total No. of Pair Mating** | 70 | 83 |
| **Fertile** | 28 (40%) | 18 (21.7%) |
| **Sterile** | 42 (60%) | 55 (66.3%) |

The similarity in the pattern of development and the defects associated with it between the *l(3)tb* trans-heterozygotes and homozygotes provide a strong genetic proof of allelism between *l(3)tb* and *DCP2*.

### *DCP2*^*l(3)tb*^ is an insertion allele of *DCP2*

Fine mapping, performed by overlapping PCR identified the region (**Supplementary Figure S5**), amplified by the primer pair, Dbo_F, and DCP2_P19_R, to span the candidate region. The region, which is of 945 bp (3L: 15826279..15827223) in the wild type and comprises of the 5’UTR coding region of *DCP2*, the adjacent intergenic region and the proximal part of the neighboring gene, *dbo*, showed absence of amplification in the DNA of *l(3)tb* homozygotes, highlighting it as the candidate lesion. Long range PCR using the same pair of primers revealed a large amplicon of ∼8.5 kb in the mutant against the 945 bp amplicon in the wild type genome, subjected to same thermal cycling parameters (**Figure 8B**). Amplification of the full length gene *DCP2* using primers residing outside the gene revealed a large amplicon of ∼17kb from the mutant genome as against the 8.6 kb (3L: 15811576..15820204) wild type amplicon (**Figure 8A**). The pGEM-T-812 probe, which corresponds to the candidate region in the wild type, hybridized with all the amplicons implying the fidelity of amplification (**Figure 8A and B**). This was further corroborated by sequencing of the amplicon terminals (data not shown). On digesting wild type and mutant genomic DNA with enzymes *Hin*dIII and *Bam*HI and subsequent hybridization with the pGEM-T-812 probe, completely different banding profiles were observed. While the *Hin*dIII digested DNA showed a band at ∼ 2.1 kb in the wild type genome, the *l(3)tb* genome showed a single band at ∼ 10 kb, the size difference being almost in agreement with the banding profile exemplified by *Bam*HI digested DNA, wherein, the wild type genome showed a band at ∼ 10.2 kb and the mutant at ∼ 18 kb (**Figure 8D**). These results imply the presence of an insertion at the candidate region in *DCP2*, in the *DCP2*^*l(3)tb*^ genome and that *DCP2*^*l(3)tb*^ is an insertion allele.

**Figure 8.**
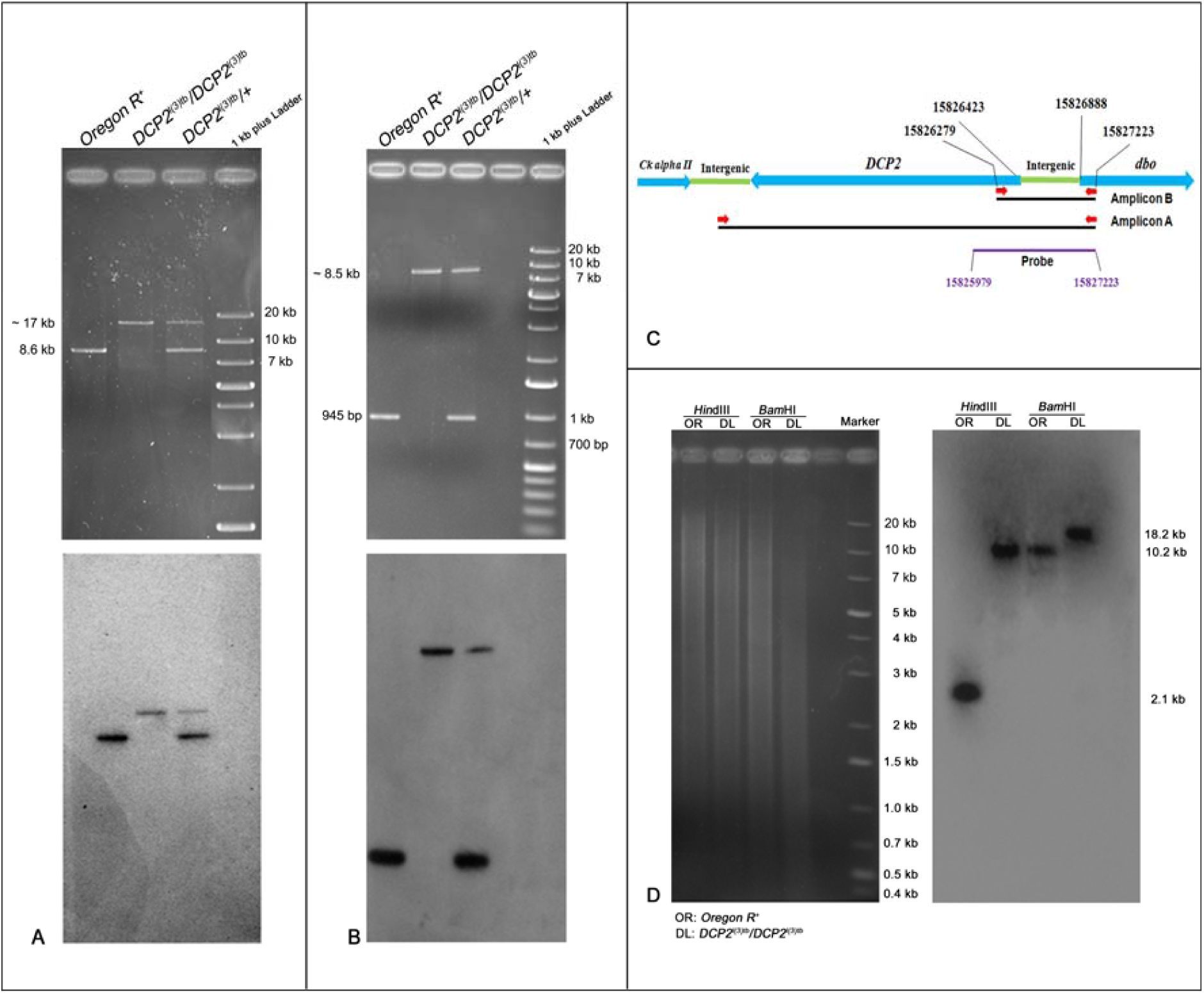
Gel electrophoretogram showing the PCR analysis of the full-length *DCP2* (A) and candidate region (B) in the wild type, mutant and the heterozygote. The schematic in C shows the gene arrangement along the chromosome along with the important coordinates. The primers are indicated by red arrows. For amplification of the full – length gene, the wild type amplicon is of 8.6 kb while the mutant amplicon is sized ∼17 kb (A, upper half), whereas the wild type amplicon for the candidate region is of 945 bp while the mutant amplicon is sized ∼8.5 kb (B, upper half). The heterozygote harbors both the alleles (wild type and mutant) and thus shows both the amplicons. The lower half in both A and B shows the the blot of the same probed with the pGEM-T-812 probe which spans the candidate mutated region in *DCP2* and is represented by the purple line. D shows the gel electrophoretogram and Southern blot of *DCP2* in the wild type and mutant genome. *Hin*dIII digested genomic DNA showed banding at ∼ 2.1 kb in the wild type genome as against ∼ 10 kb in the mutant genome, the size difference being almost in agreement with the banding profile exemplified by *Bam*HI digestion, with the wild type genome hybridizing at ∼ 10.2 kb and the mutant at ∼ 18 kb.

### The *DCP2*^*l(3)tb*^ genome harbors *Gypsy*-LTR like sequence in 5’UTR coding region of *DCP2* and expansion of adjacent upstream intergenic AT-rich sequence

In order to identify the functional genomics of mutations, it is essential to deduce the nucleotide sequence of the mutation, and thus, a convergent bi-directional primer-walk was initiated with the primer pair which identified the presence of insertion in *DCP2* in the *l(3)tb* genome. On sequencing, the *DCP2*-proximal end showed presence of wild type sequence till 3L: 15826410 after which a 444 bp AT-rich sequence was detected (**Supplementary Figure S6 B-1**), which did not show any resemblance with the wild type sequence present at the region whereas the *dbo*-proximal end showed complete wild type sequence profile (3L: 15827143..15826738) (**Supplementary Figure S6 B-2**). On homology search to identify the novel sequence obtained, the sequence showed homology with the *Gypsy* LTR sequence of *Drosophila*. On searching for *DCP2* promoters in the Eukaryotic Promoter Database, SIB and aligning the sequence coordinates of the 444 bp insertion, it was found that the insertion is downstream to the transcription start site (TSS) of *DCP2*, which is at 3L: 15826420. On designing a new pair of primers from the distal part of the reads obtained above, long range PCR was first performed with *DCP2*^*l(3)tb*^ and wild type genomic DNA. Although no amplification was observed with the wild type DNA, the *DCP2*^*l(3)tb*^ DNA showed amplicons of sizes ∼7.2 kb, ∼3 kb, ∼2.8 kb and ∼430 bp with the 430 bp amplicon showing the highest concentration as observed from the electrophoretogram (**Figure 9B**). This amplification profile resembled that of tandem repeat bearing regions. To confirm the repetitive nature of the sequence, Southern hybridisation was performed with the same electrophoretogram. The 430 bp amplicon was eluted from the gel, cloned in pGEM-T vector and used as a probe. The probe showed complete hybridization with all of the amplicons indicating the repetitive nature of the sequence present downstream (**Figure 9B**). Sequencing of the 430 bp amplicon revealed an AT-rich sequence. Homology search identified the sequence to be homologous to the distal part of the *DCP2* UTR and the adjacent intergenic sequence between *DCP2* and *dbo*, the coordinates being 3L: 15826407..15826716. After aligning the present set of reads with the previous set, a sequence duplication was observed for 5’-T-A-T-A-3’, flanking the *Gypsy*-LTR insertion (**Supplementary Figure S6 B-3 and 4**). The present set of sequencing reads also confirmed that the LTR insertion (3L: 15826407..15826407) was indeed prior to the completion of the UTR (3L: 15826423). Copy number variation analyses of the intergenic sequence *vs.* the internal control through PCR in the wild type and the mutant *DCP2*^*l(3)tb*^ showed a sharp increase in the amplicon concentration of the intergenic sequence in *DCP2*^*l(3)tb*^ against the internal control as evidenced from the gel electrophoretogram (**Figure 9C**). Comparison of the fluorescent intensity of the bands (intra and inter-genotype) showed relatively high ratio of concentrations of the amplicon to the internal control (*Dsor*) amplicon as observed from the graphical analyses (**Figure 9D**). On the basis of the results obtained from rough and fine mapping, the architecture of the mutant allele, *DCP2*^*l(3)tb*^ is depicted in **Figure 9E**, which shows the bipartite nature of the mutation, *viz.*, amplification of the intergenic sequence between *DCP2* and *Dbo* as well as an insertion of 444 bp *Gypsy* LTR-like sequence immediately downstream to the TSS of *DCP2*.

**Figure 9.**
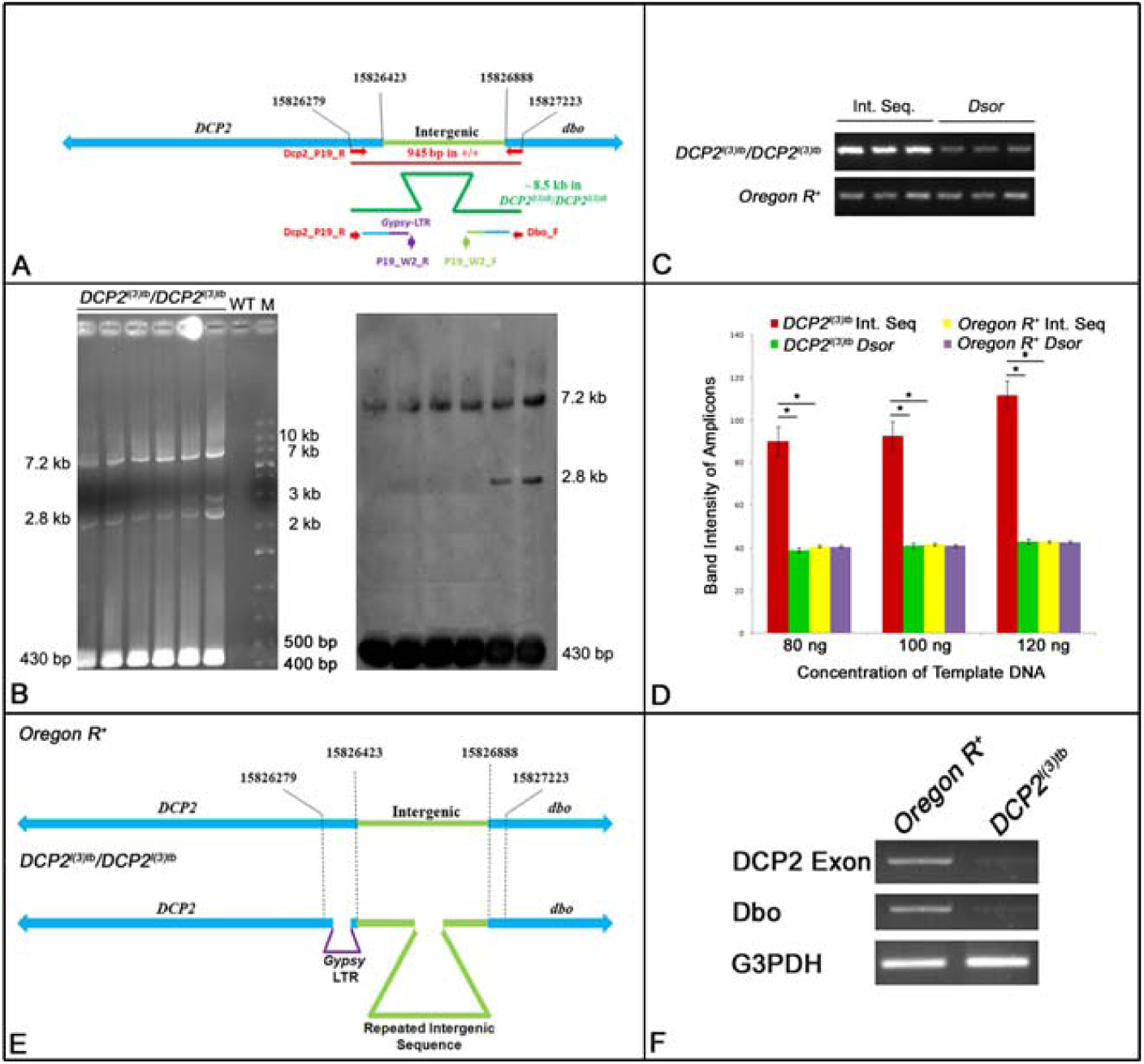
Gel electrophoretogram showing PCR amplification profile obtained by the primers used (P19_W2_R and P19_W2_F) for the “second step” of walking (B). The alignments of the sequences uncovered in the “first step” are shown as thin lines colored as per homology with the wild type sequence. The region amplified here lies subsequent to the sequence uncovered by the initial primers (A; Dbo_F and Dcp2_P19_R, shown in red arrows). Mentioned alongside the electrophoretogram are the semi-logarithmic estimates of the amplicon size. Shown alongside is the blot of the same hybridized with the probe generated from the ∼430 bp amplicon. Semi-quantitative PCR to detect change in copy number of the intergenic region in the *DCP2*^*l(3)tb*^ genome (C) shows increased amplification of the intergenic sequence in *DCP2*^*l(3)tb*^ genome as compared to the *Dsor* (control) amplicons in both the genomes. Shown in D is a histogram comparing the fluorescence intensity of PCR amplicons obtained from amplification of the intergenic sequence and the control sequence from the *DCP2*^*l(3)tb*^ genome and the wild type genome. The schematic in E shows the architecture of the mutant allele, *DCP2*^*l(3)tb*^ based on the results obtained from fine mapping. Semi-quantitative RT-PCR analyses of transcription from *DCP2* and *Dbo* in the wild type and *DCP2*^*l(3)tb*^ homozygotes (F) shows decreased titre of mRNA from both genes in the tumorous individuals.

### *DCP2*^*l(3)tb*^ is a *DCP2* hypomorph alongwith low expression of the neighbouring gene, *Dbo*

Following the identification of the genomic architecture of the allele, it was imperative to determine the expression potential of the allele. Semi-quantitative RT-PCR analyses confirmed the hypomorphic nature of the allele wherein the mutant showed extremely low levels of expression of *DCP2* (**Figure 9F**). Since the intergenic region between *DCP2* and *Dbo* is upstream to either and bears an expansion, *Dbo* transcript titres were also examined wherein they showed extremely lowered expression (**Figure 9F**). At present, it is doubtful whether the lowered Dbo level is a cause or an effect of the mutation, since Dbo (Smac/Diablo/Henji) is a pro-apoptogenic molecule which the inhibitor of apoptotic proteins (IAP), and its lowered levels therefore serve as a prognostic marker of tumor progression in human carcinomas (Martinez-Ruiz et al, 2008). Again, *Dbo* expresses strongly in the neuronal tissues at the synapse (Wang et al, 2016) and its perturbation causes alteration in neuro-muscular function. Being a multi-faceted molecule, altered expression of the same may have some contribution to the tumourigenesis since the tumor primarily affects brain which is an integral part of the CNS.

### Tumor caused by *DCP2* is hyperplastic with elevated Cyclins A and E

Since the mutation showed all the hallmarks of classical tumor suppressors (Merz et. al., 1990), we endeavored to characterize the perturbations in cellular physiology caused in the wake of tumourigenesis. RT-PCR analyses depicted elevated levels of Cyclins E (G1/S phase cyclin) and A (G2/M phase cyclin), which are indicative of increased cell proliferation and rapid cell cycles (**Figure 10C**). Immunolocalisation studies confirmed the elevated expression of Cyclin E as well (**Figure 10A and B**). On observing closely, the regular arrangement of cells in the brain hemisphere and optic lobes in the wild type is severely disrupted in the mutant along with superfluous growth and increased number of mitotic nuclei. The enlarged brain lobes, increased number of mitotic nuclei and disruption of the regular arrangement of cells in the mutant, concomitant with elevated expression of cyclins A and E clearly imply the tumourous nature of the mutant.

**Figure 10.**
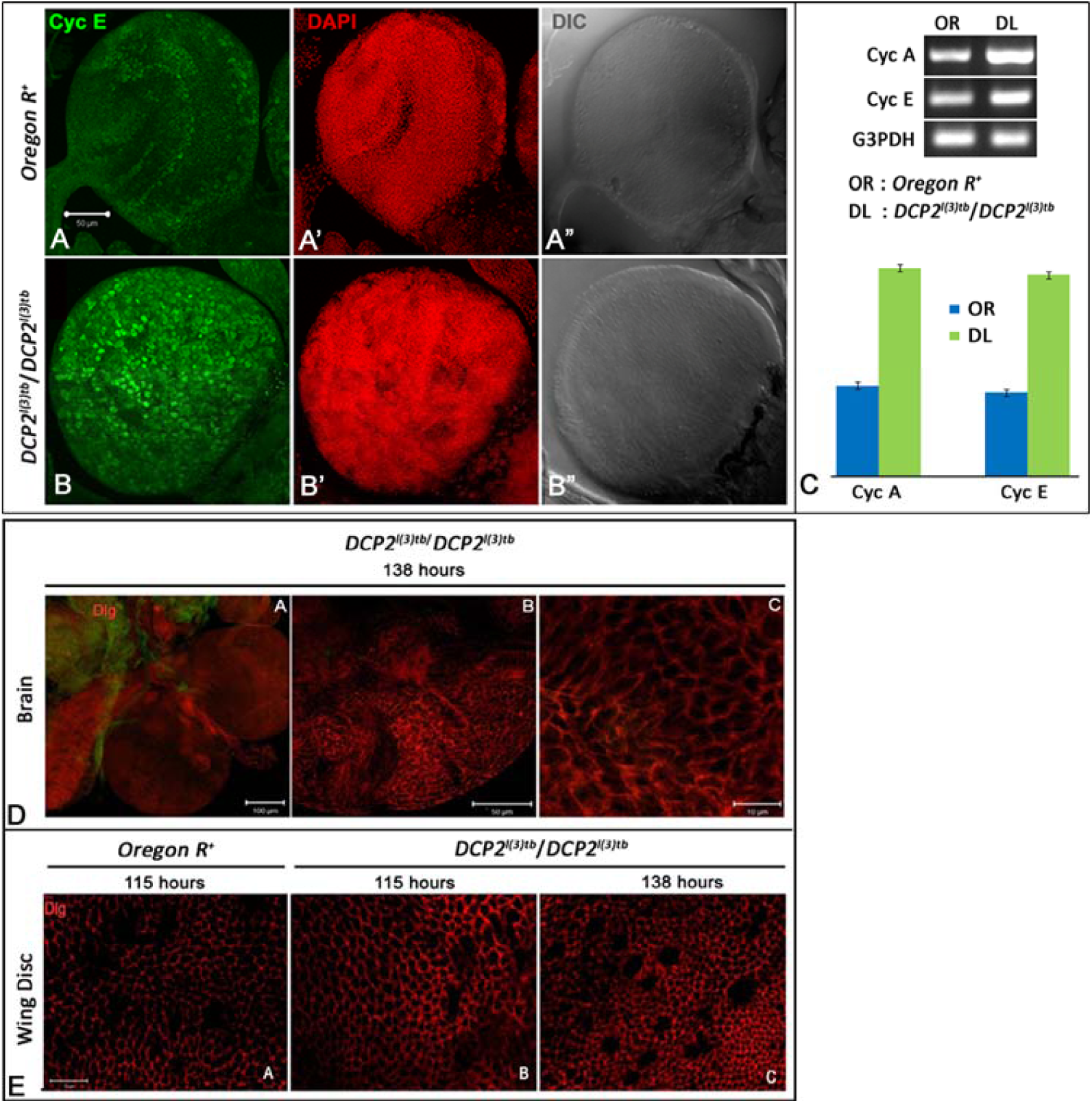
Immunolocalisation of Cyclin E shows elevated expression in the tumorous larval brains of *DCP2*^*l(3)tb*^ homozygotes (B) as compared to the wild type (A). Semi-quantitative analyses of mRNA expression of Cyclins A and E show similar elevation in the brain of *DCP2*^*l(3)tb*^ homozygotes (C). Expression of Discs-large in the brain (D) and wing discs (E) of the tumorous individuals did not show appreciable loss. At 138h AEL, the wing discs showed increase in cell number concomitant with decrease in cell size (E.C) whereas, at the same stage, the tumorous brain shows increased number of cells at in the optic lobe (D).

When the tumorous brains (**Figure 10D**) and wing discs (**Figure 10E**) were examined for the expression of the polarity marker Discs large (Dlg), both tissues did not show appreciable loss of polarity. On a closer look, the wing discs at 138h AEL showed increase in cell number concomitant with decrease in cell size (**Figure 10E-C**). At the same stage, the tumorous brain shows increased number of cells at in the optic lobe (**Figure 10D**). Usually, neoplastic tumours are metastatic and the tumour cells lose their polarity to acquire the mesenchymal-like fate, delaminate from the matrix and migrate (Miles et al, 2011). In contrast, hyperplastic tumours do not show appreciable loss of polarity even in later stages of tumourigenesis, since in these tumours the cells do not delaminate, but remain adhered to the original tissue matrix, but keep on dividing. The expression pattern of Dlg shows retention of polarity even at 138h of development, which is an extremely late and delayed 3^rd^ instar larval stage, which implies the tumour to be an over-proliferative, hyperplastic one. Again, this is well in agreement with the Cyclin E staining pattern, and taken together, they imply increased cell division, which essentially requires increased and rapid cell cycles.

### Global overexpression of *DCP2* rescues mutant phenotypes associated with *l(3)tb*

Global over-expression of *DCP2* using ubiquitous GAL4 drivers (*Act5C-GAL4* or *Tub-GAL4*) in the mutant homozygous *l(3)tb* individuals rescued the larval and pupal lethality. **Table 6** shows the genotype and fate of the progeny as scored from the rescue experiment. As can be seen, for over-expression of *DCP2* using *Act5C-GAL4*, out of 35.1% (N=155) non-tubby progeny (*l(3)tb* homozygous background), *i.e., Act5C-GAL4/CyO* or *Sp; l(3)tb:UAS-DCP2/l(3)tb*, 21.3% (N=94) and 13.8% (N=61) segregated as curly (*Act5C-GAL4/CyO; l(3)tb:UAS-DCP2/l(3)tb*) and non-curly (or with sternopleural bristles: *Act5C-GAL4/Sp; l(3)tb:UAS-DCP2/l(3)tb*), respectively. Similarly, while over-expressing using *Tub-GAL4*, we obtained 37% (N=166) non-tubby progeny, *i.e., UAS-DCP2/CyO* or *Sp*; *l(3)tb:Tub-GAL4/l(3)tb*, out of which, 17.2% (N=77) were curly (*UAS-DCP2/CyO*; *l(3)tb:Tub-GAL4/l(3)tb*) while 19.8% (N=89) were non-curly (*UAS-DCP2/ Sp*; *l(3)tb:Tub-GAL4/l(3)tb*).

**Table 6.** Global overexpression of *DCP2* rescues the mutant phenotypes exhibited by *l(3)tb* homozygoytes.

| Genetic Crosses |  | *Act5C-GAL4/CyO*;<br>+/+ |  | +/+; *Tub-GAL4/TM6B* |  | *Act5C-GAL4/CyO*;<br>*l(3)tb/TM6B* |  | *UAS-DCP2/CyO*;<br>*l(3)tb/TM6B* |  |
| --- | --- | --- | --- | --- | --- | --- | --- | --- | --- |
|  |  | *Act5C-GAL4/CyO*;<br>+/+ |  | +/+; *Tub-GAL4/TM6B* |  | *Sp/CyO*; *l(3)tb*:<br>*UAS-DCP2/TM6B* |  | *Sp/CyO*; *l(3)tb*:<br>*Tub GAL4/TM6B* |  |
|  |  | Homozygotes die as embryos or early larvae |  | Homozygotes die as embryos or early larvae |  | *CyO* and *TM6B* homozygotes die as embryos or early larvae |  | *CyO* and *TM6B* homozygotes die as embryos or early larvae |  |
| 01. | Eggs | **750** |  | **790** |  | **1050** |  | **1245** |  |
| 02. | Unfertilised Eggs | 39 | 5.2% | 37 | 4.7% | 68 | 6.5% | 86 | 6.9% |
| 03. | Fertilised Eggs | 711 | 94.8% | 753 | 95.3% | 982 | 93.5% | 1159 | 93.1% |
| 04. | Dead Embryos | 304 | 42.8% | 357 | 47.4% | 434 | 44.2% | 525 | 45.3% |
| 05. | Dead 1<sup>st</sup> and 2<sup>nd</sup> instar Larvae | 8 | 1.12% | 19 | 2.52% | 34 | 3.46% | 57 | 4.92% |
| 06. | Dead 3<sup>rd</sup> instar larvae | 2 | 0.28% | 5 | 0.66% | 72 | 7.33% | 128 | 11.04% |
| 07. | Pupae | **397** | 55.8% | **372** | 49.4% | **442** | 45.0% | **449** | 38.7% |
| 08. | Dead Pupae | 17 | 4.3% | 11 | 2.9% | 21 | 4.8% | 23 | 5.1% |
| 09. | Eclosion following over-expression of *DCP2* in homozygous *l(3)tb* background | - | - | - | - | *Act5C-GAL4/CyO*;<br>*l(3)tb*:*UAS-DCP2/l(3)tb* |  | *UAS-DCP2/CyO*;<br>*l(3)tb*: *Tub-GAL4/l(3)tb* |  |
|  |  |  |  |  |  | 94 | **21.3%** | 77 | **17.2%** |
|  |  | - | - | - | - | *Act5C-GAL4/Sp*;<br>*l(3)tb*:*UAS-DCP2/l(3)tb* |  | *UAS-DCP2/Sp*;<br>*l(3)tb*: *Tub-GAL4/l(3)tb* |  |
|  |  |  |  |  |  | 61 | **13.8%** | 89 | **19.8%** |

In both the cases of overexpression, all non-tubby progeny pupated, devoid of any developmental anomalies reminiscent of *l(3)tb* mutation and emerged as flies. Thus, the rescue of the mutant phenotypes observed in *l(3)tb* homozygotes by global overexpression of *DCP2* iteratively substantiates the fact the *l(3)tb* is an allele of *DCP2* and that the tumour is caused solely owing to the loss of expression of *DCP2*.

## Summary and Conclusion

In *Drosophila*, DCP2 is the only decapping enzyme present and thus is extremely important for a number of growth processes throughout development. In other organisms as well, it is well conserved and has fundamentally important roles in development (Xu et al, 2006; Ma et al, 2013), DNA replication (Mullen and Marzluff, 2008; Schmidt et al, 2011), stress response (Hilgers et al, 2006; Xu and Chua, 2012), synapse plasticity (Hillebrand et al, 2010), retrotransposition (Dutko et al, 2010) and viral replication (Hopkins et al, 2013). In *Arabidopsis, DCP2* loss-of-function alleles show accumulation of capped mRNA intermediates, lethality of seedlings and defects in post-embryonic development, with no leaves, stunted roots with swollen root hairs, chlorotic cotyledons and swollen hypocotyls (Goeres et al, 2007; Iwasaki et al, 2007; Xu et al, 2006). In humans as well, chromosomal deletions of 5q21-22, the region harboring *DCP2* is frequently observed in lung cancers (Hosoe et al, 1994; Mendes-da-Silva et al, 2000), colorectal cancer (Delattre et al, 1989) and oral squamous cell carcinoma (Mao et al, 1998). Hence, *DCP2* has an unexplored role in development and/or cell cycle progression across phyla, which needs to be investigated. Since the physiology of an organism is tightly regulated by the optimized titres of gene expression programs, a global loss of *DCP2* may lead to perturbed mRNA titres which in turn may alter the cellular response to such dismal conditions and eventually lead to drastic physiological disorders such as tumourigenesis. Although we are unsure of the exact mechanism(s) by which a loss of *DCP2* leads to tumourigenesis, our findings in the novel allele, *DCP2*^*l(3)tb*^, propose an absolutely novel role of *DCP2* in tumourigenesis and identify *DCP2* as a candidate for future explorations of tumourigenesis.

## Acknowledgements

The authors acknowledge Prof. L. S. Shashidhara, IISER Pune, India and Prof. Volker Hartenstein, University of California, USA for sharing the Anti-Armadillo and Anti-Deadpan antibodies, respectively. We sincerely acknowledge Prof. Rajiva Raman and Dr. Rachana Nagar of our lab for assistance in fine mapping of the mutation and in analyses and interpretation of the results obtained therein. We duly acknowledge the National Facility for Laser Scanning Confocal Microscopy, UGC-CAS, DST-FIST of Department of Zoology, Banaras Hindu University and DST-PURSE, UGC-UPE to the Institute. We sincerely thank Department of Science and Technology (DST) for financial support to JKR.

## Ethical Approval

All studies were performed as per ethical guidelines. All applicable international, national and institutional guidelines for the care and use of *Drosophila* were followed.

## Supplementary Figures

**Figure S1.**
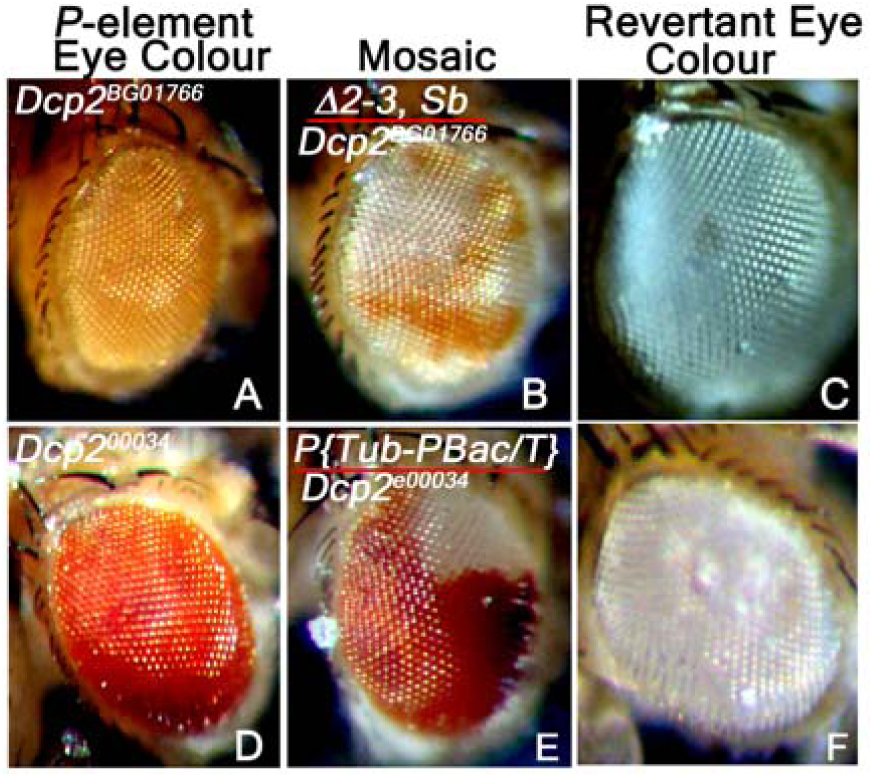
Reversion analysis by the excision of *piggyBac* transposon in *DCP2*^*e00034*^ with the help of *piggyBac* specific transposase source, *CyO, P{Tub-Pbac}2/Wg*^*SP-1*^ and similarly by the excision of *P*-element in *DCP2*^*BG01766*^ strain using Δ*2-3,Sb/TM6B, Tb*^*1*^, *Hu, e*^*1*^ transposase source as ‘jumpstarter stock’. DCP2 revertant white eyed F2 flies were crossed to *l(3)tb* and lethal progenies scored.

**Figure S2.**
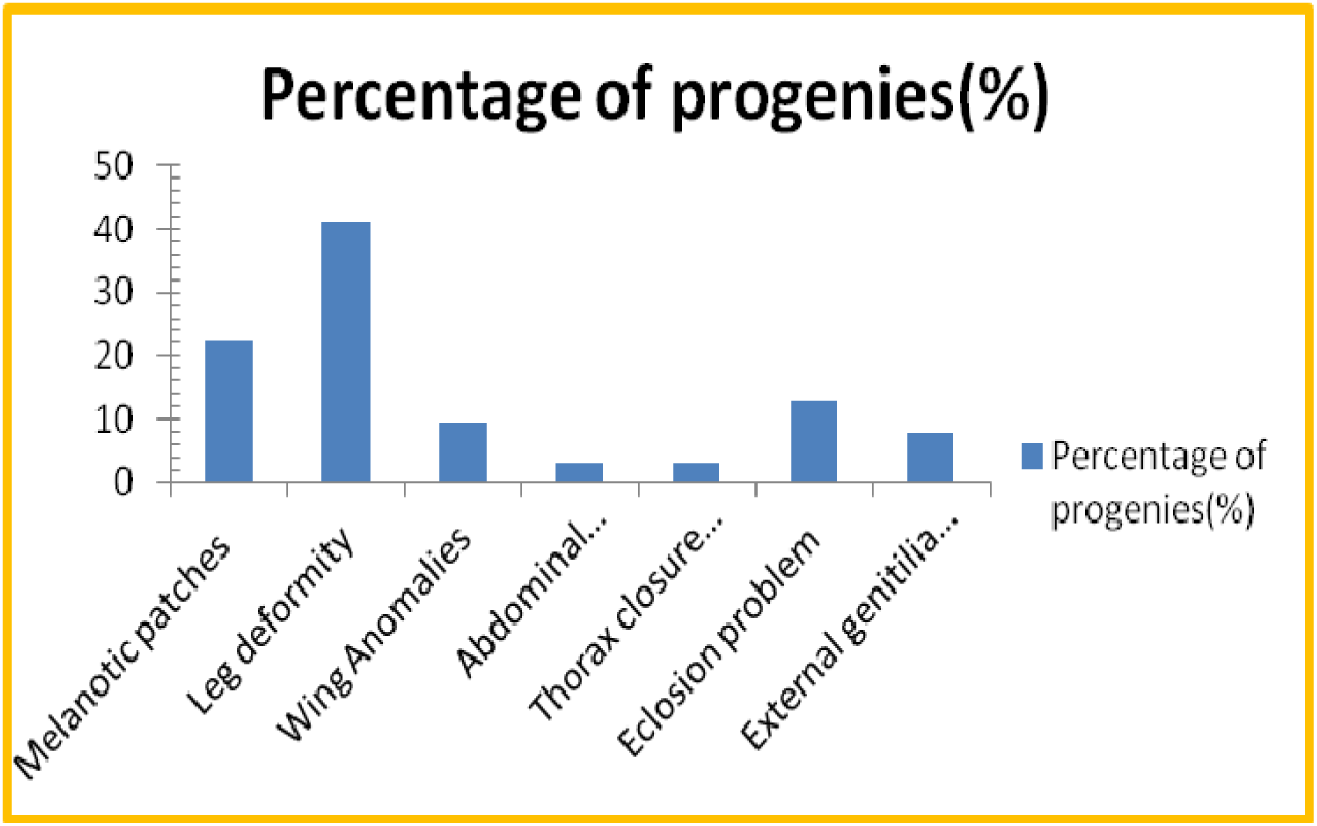
Morphological defects exhibited by escapees of adult fly trans-heterozygous for *P{GT1}DCP2*^*BG01766*^*/l(3)tb*. The phenotype includes melanotic patches (22.2%) on the cuticular exoskeleton, abnormalities in leg (41.3%), wing (10%), abdomen (3.2%) and thorax (3.2%). Many of the trans-heterozygous progeny was observed to have eclosion problem (12.7%) and males have abnormal genitalia (9.7%).

**Figure S3.**
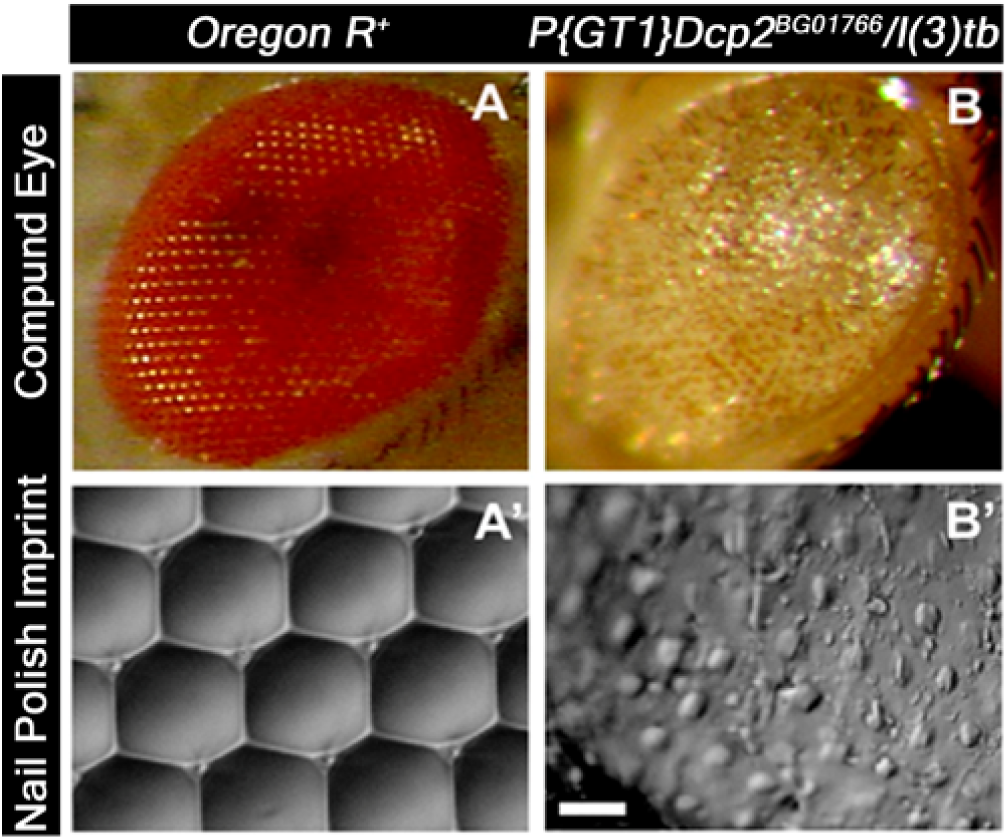
**Pronouncement of severe defects in compound eyes of the escapees having heterozygous genetic background of the mutant *l(3)tb* with lethal *P*-insertion allele *DCP2*^*BG01766*^.** Images in A and B showing the compound eye of wild type and tans-heterozygote respectively while A’ and B’ are their respective nail-polish imprint of the compound eye, viewed with the help of DIC or Nomarski microscope. The exact geometrical arrangement of ommatidia in a hexagonal pattern having each ommatidium surrounded by bristle was completely disrupted in the trans-heterozygote exhibiting the complete loss of arrangement in the ommatidial pattern. This represents the severe loss of polarity as it cues a complete disassembly of compound eye as whole. Bar represents 20µm.

**Figure S4.**
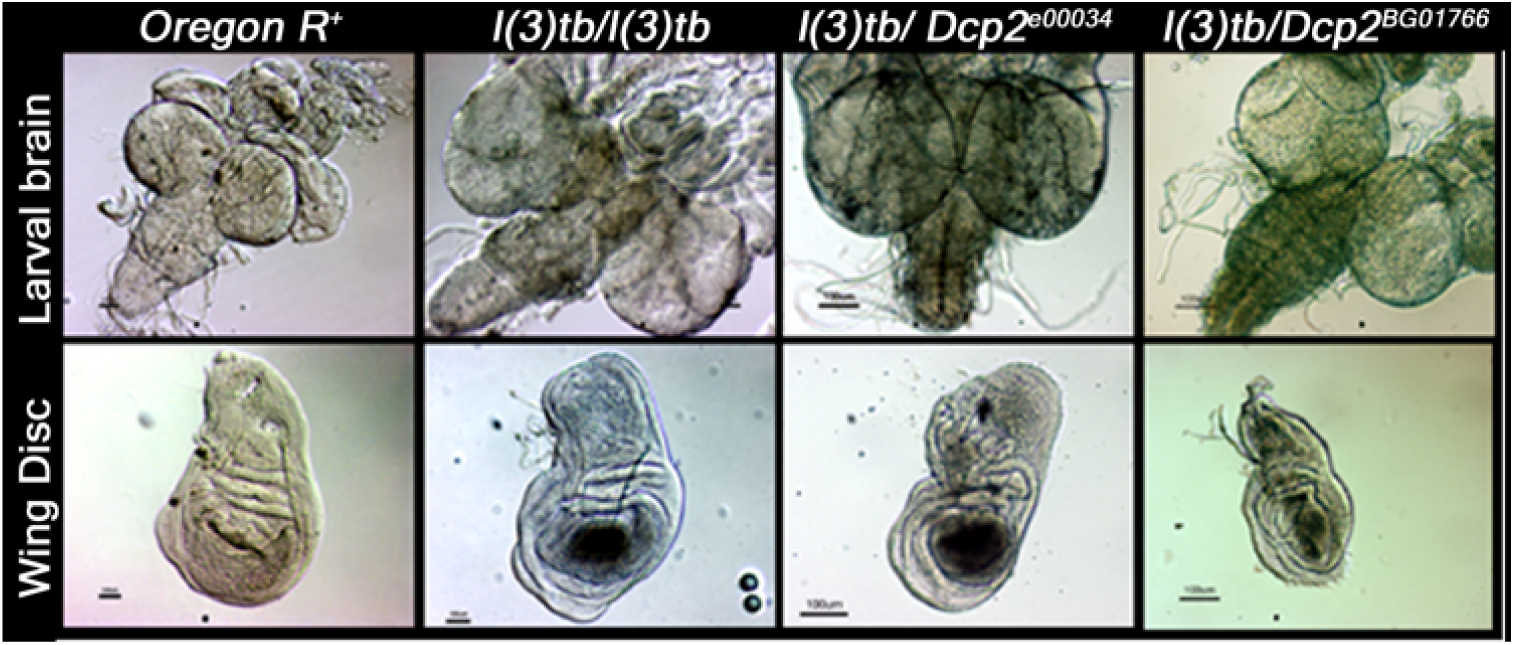
Tumorous phenotype observed in larval brain and wing imaginal discs in trans-heterozygotes *l(3)tb /PBac{RB}DCP2*^*e00034*^ and *l(3)tb /P{GT1}DCP2*^*BG01766*^ as homozygous *l(3)tb* Scale bar is 100µm.

**Figure S5.**
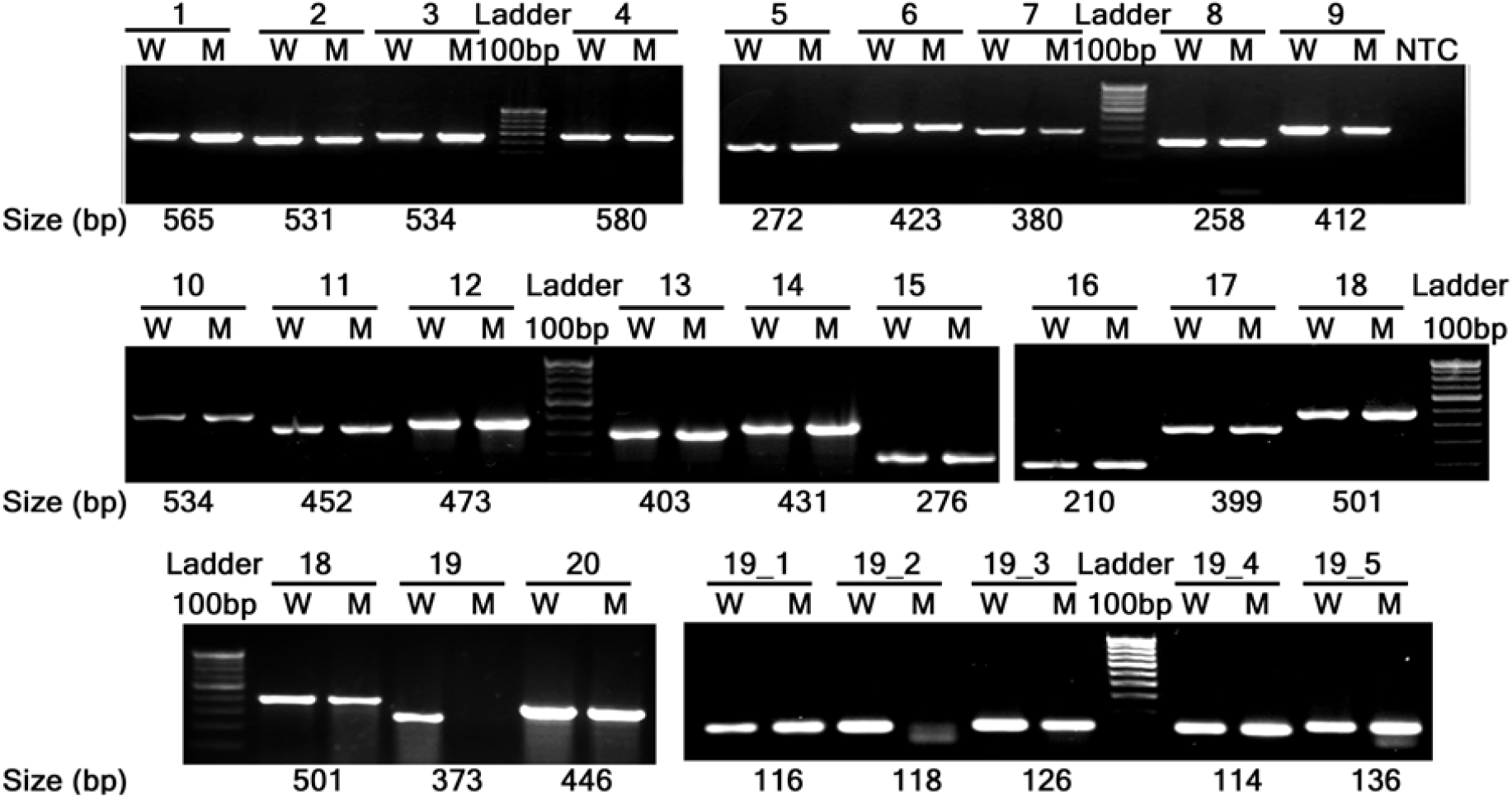
Amplification of *DCP2* using overlapping primers. All primers amplify same size of amplicon with DNA from wild type and homozygous l(3)tb mutant, except DCP2_P19 (3L:15819379..15819751) and DCP2_P19_2 (3L:15819452..15819569). This implies the probable mutation in the region.

**Figure S6.**
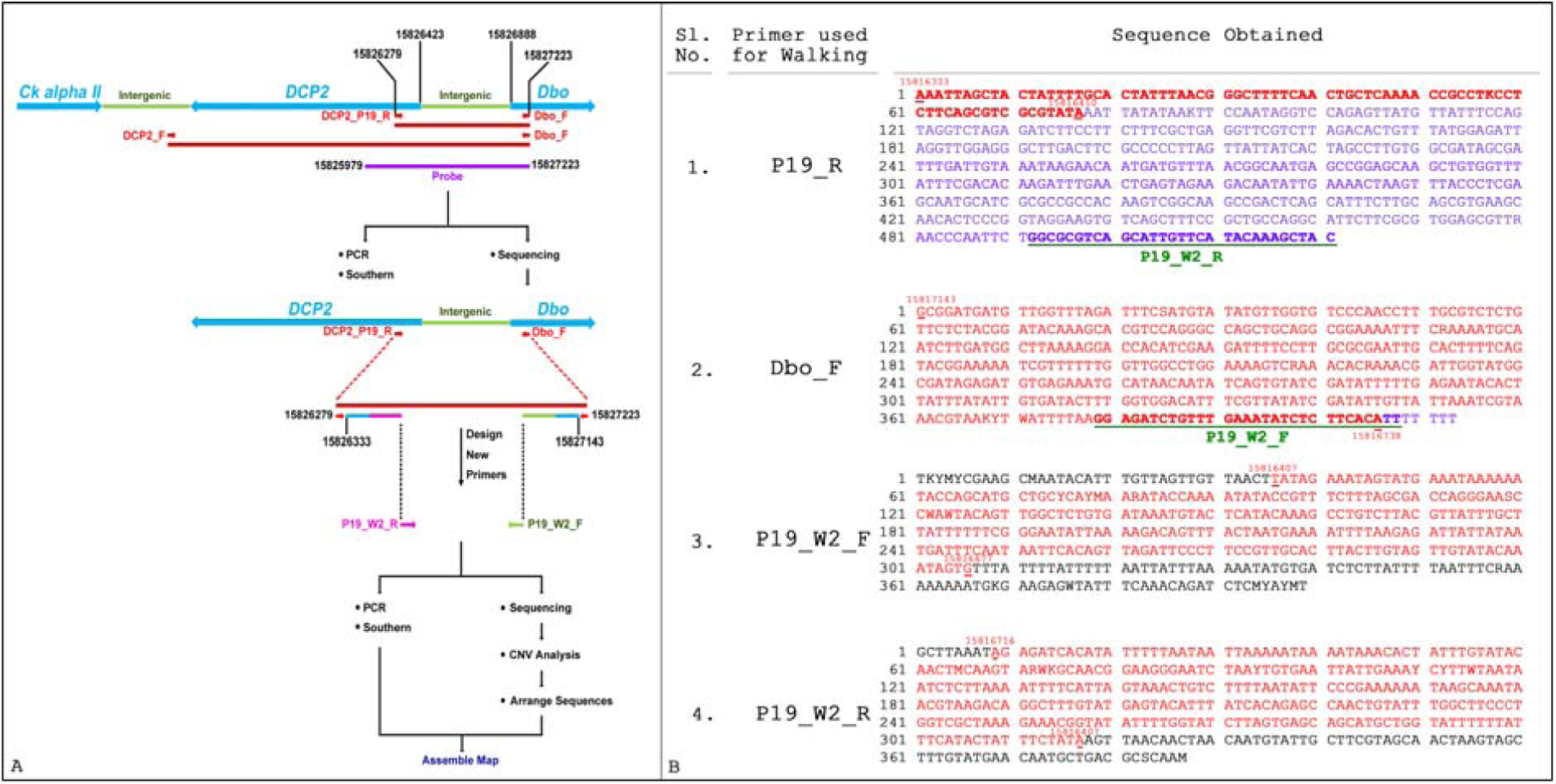
Schematic representation of the convergent bidirectional primer walking adopted for sequencing and alignment of the large amplicon obtained at the candidate region in *DCP2*^*l(3)tb*^ homozygotes (A). Shown in differently colored arrows are the primers used for sequencing during walking. Reads aligning to the gene regions are represented by blue lines, while those aligning to the intergenic regions are depicted by green lines. The primers designed are represented in similar colors depending on their alignment in the sequence. The reads obtained on sequencing with each of the four primers is shown in B. The *Gypsy*-LTR sequence is shown in purple. Underlined in 1 and 2 are the sequences used as primers for the second-step of primer walking.

## Supplementary Tables

**Table S1.**
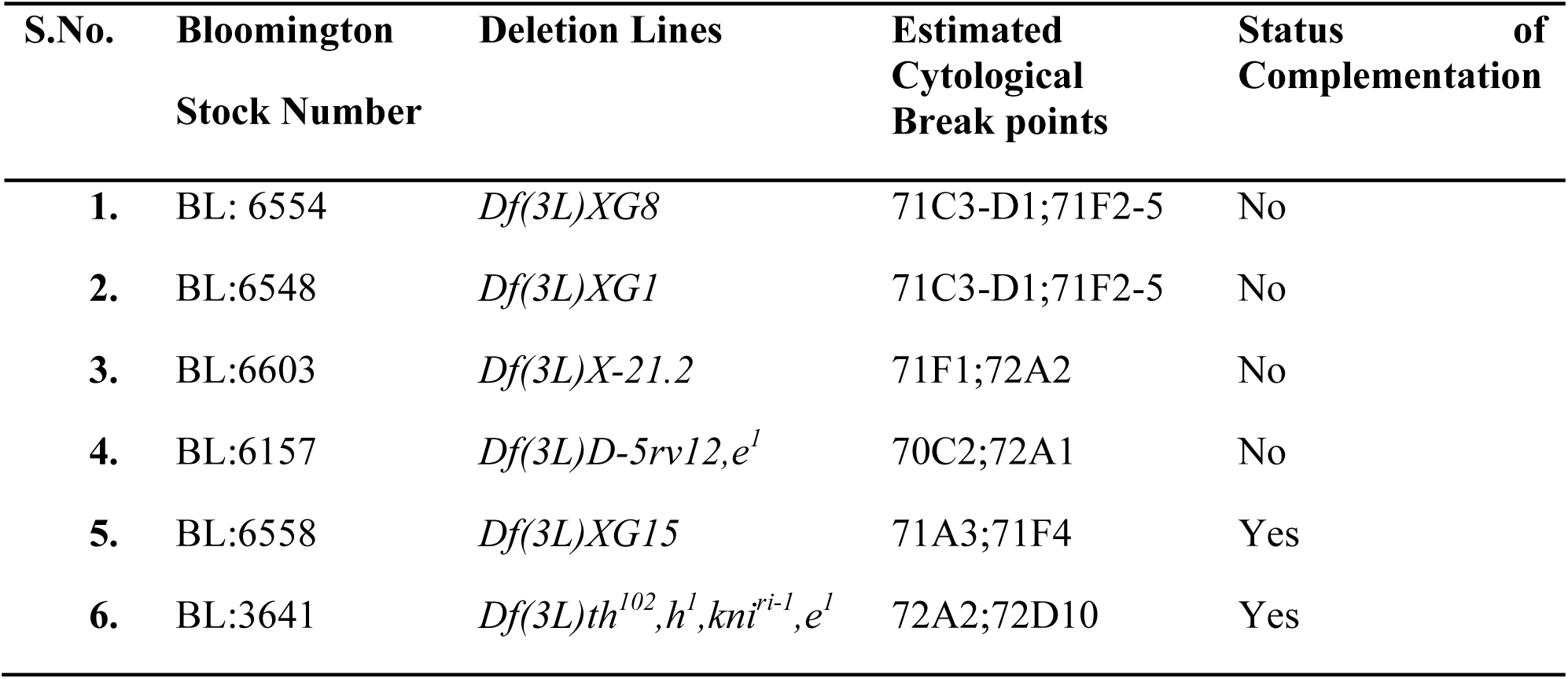
Complementation status of *l(3)tb* with cytologically mapped deletion lines.

**Table S2.**
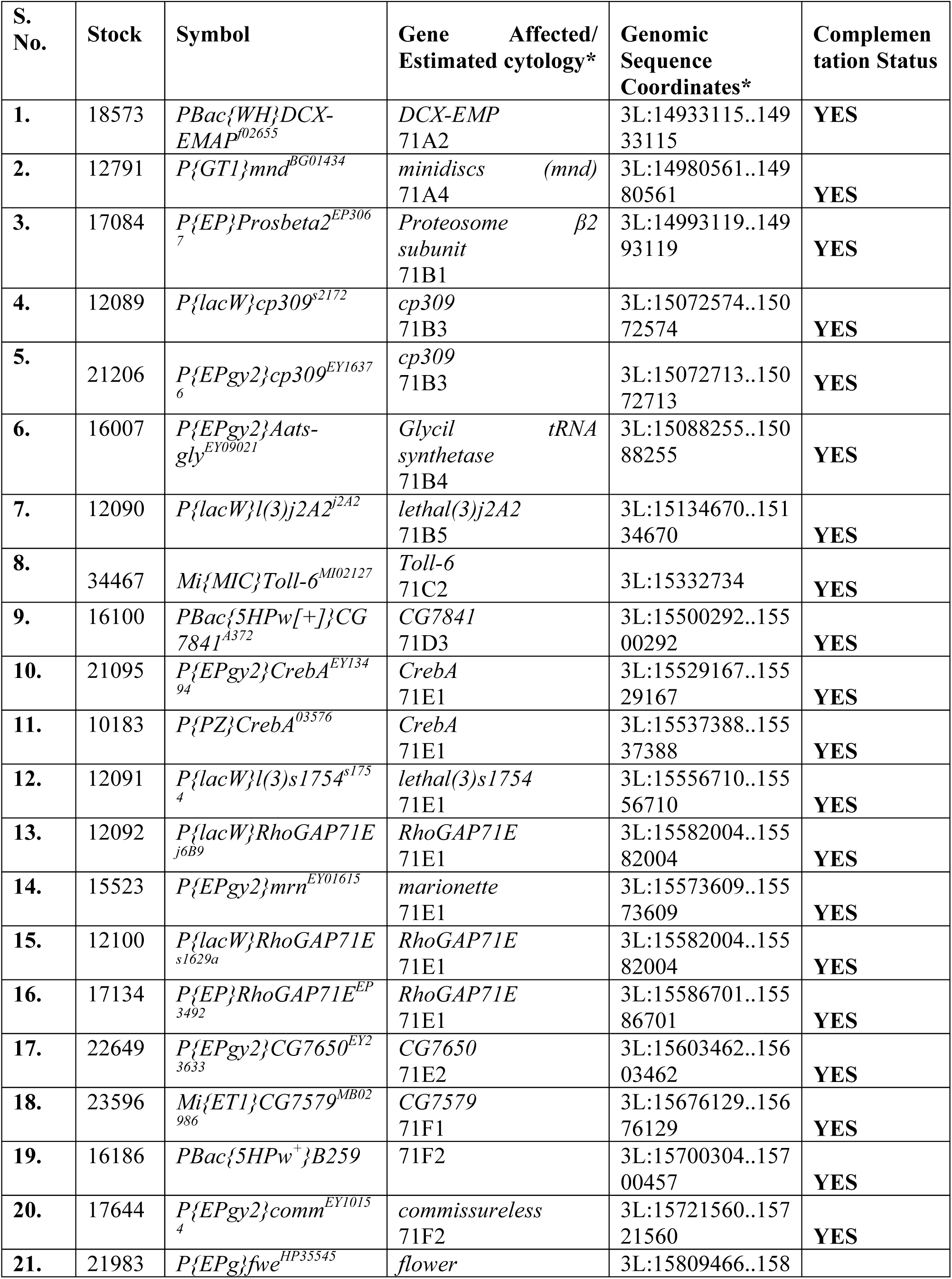
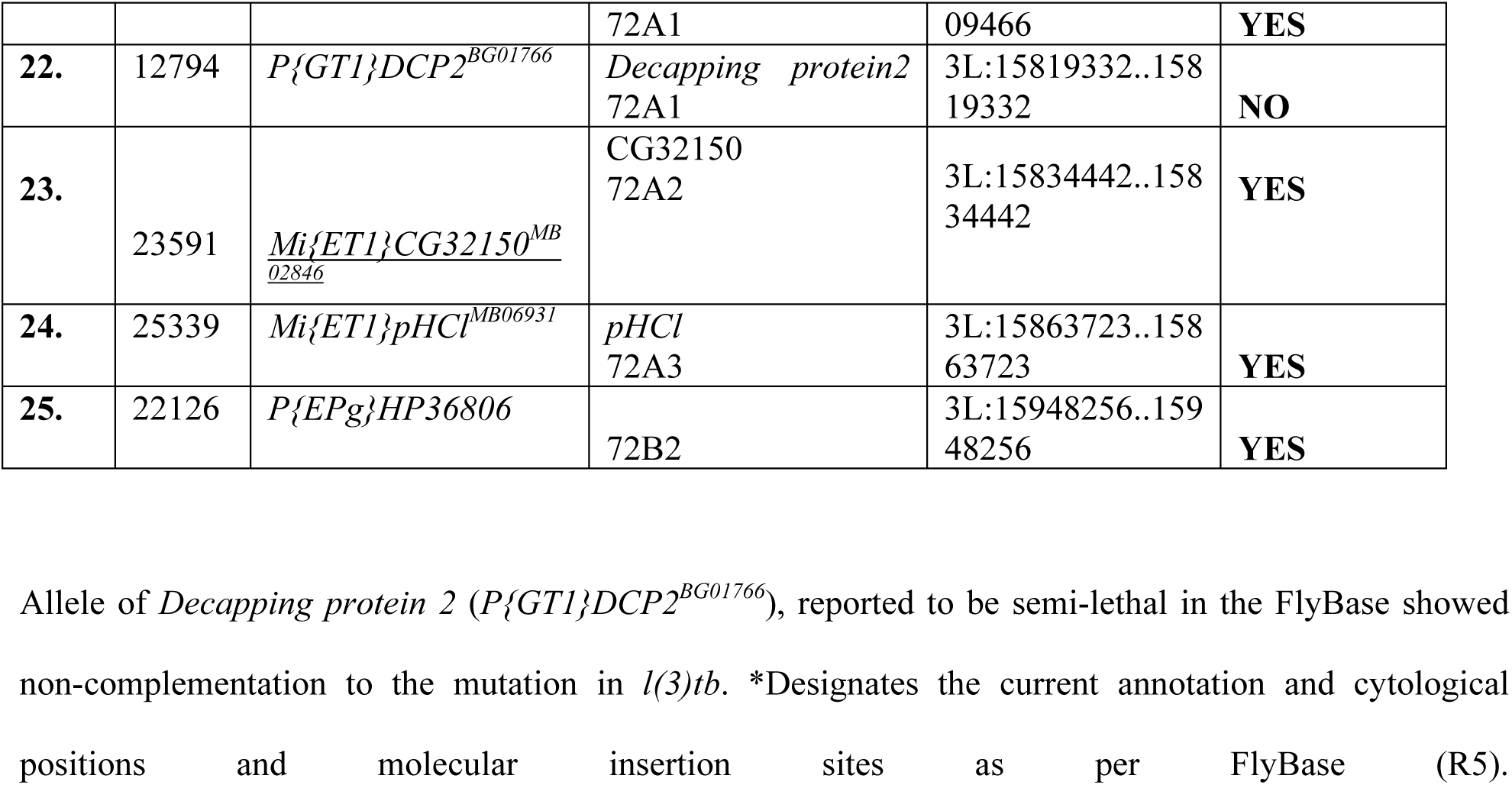
Complementation analysis of *l(3)tb* with lethal transposon insertion lines.

**Table S3.**
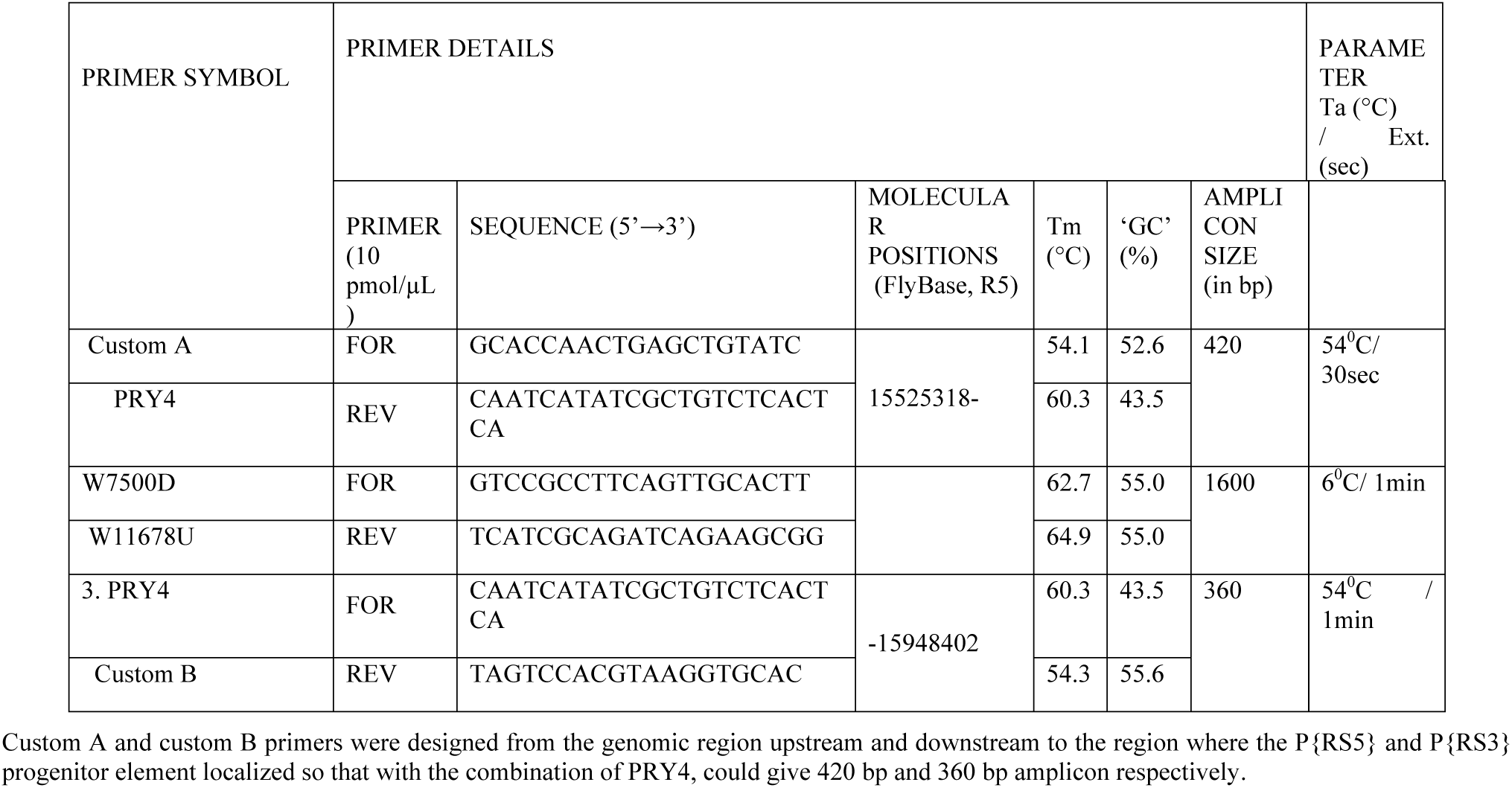
Primers used for characterizing deletion in *Df(3L)RM95*.

**Table S4.**
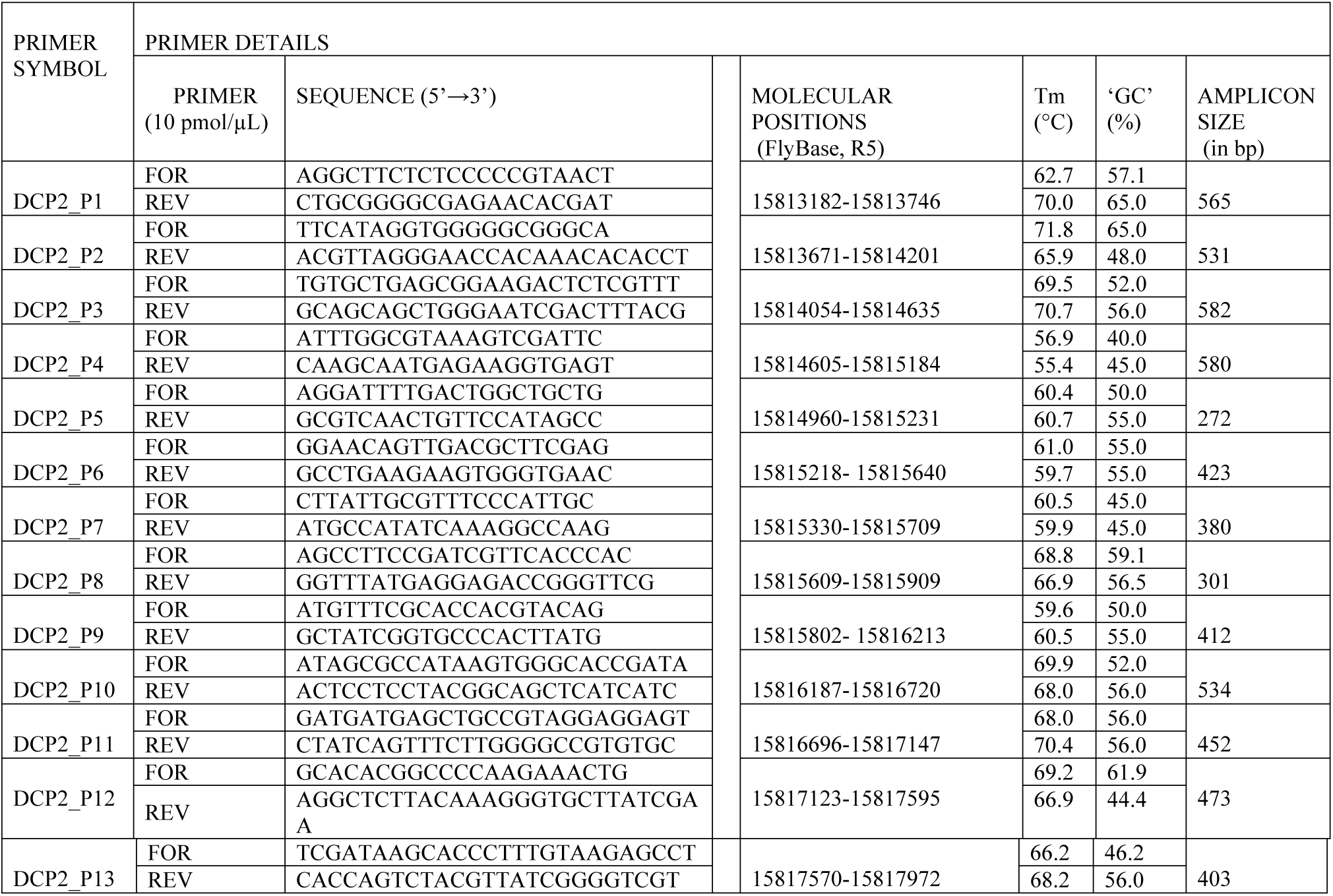
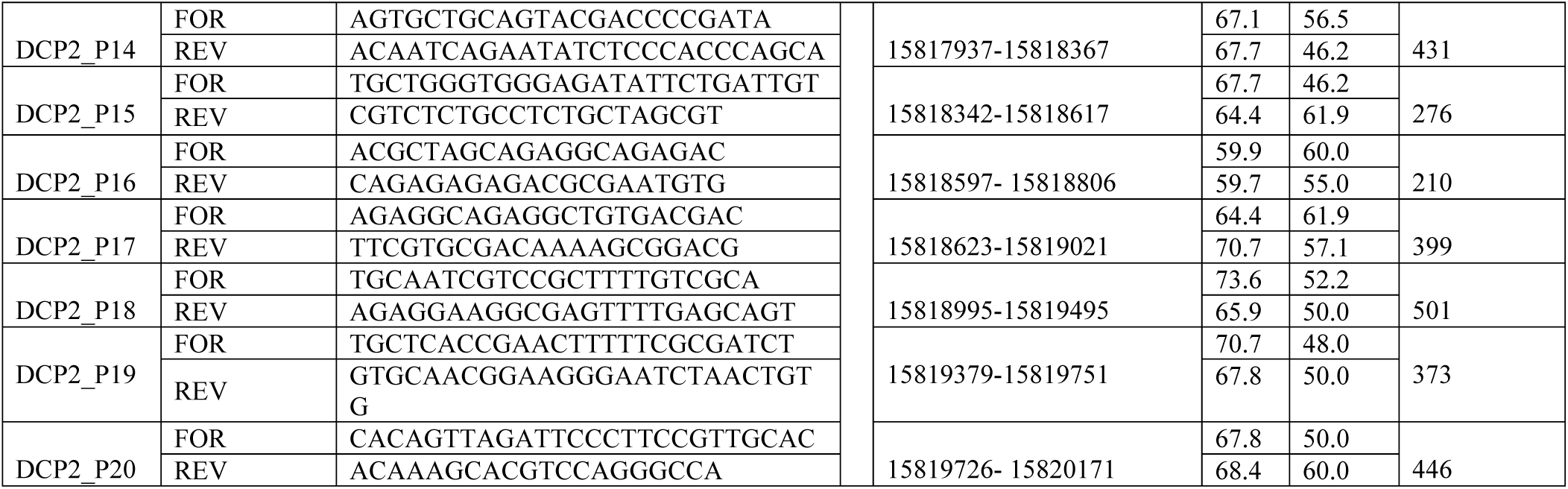
Overlapping set of primers for *DCP2* gene and thermal cycler conditions of annealing temperature and extension time for each primer pair to amplify the genomic region of *DCP2* gene in the homozygous *l(3)tb* mutant.

**Table S5.**
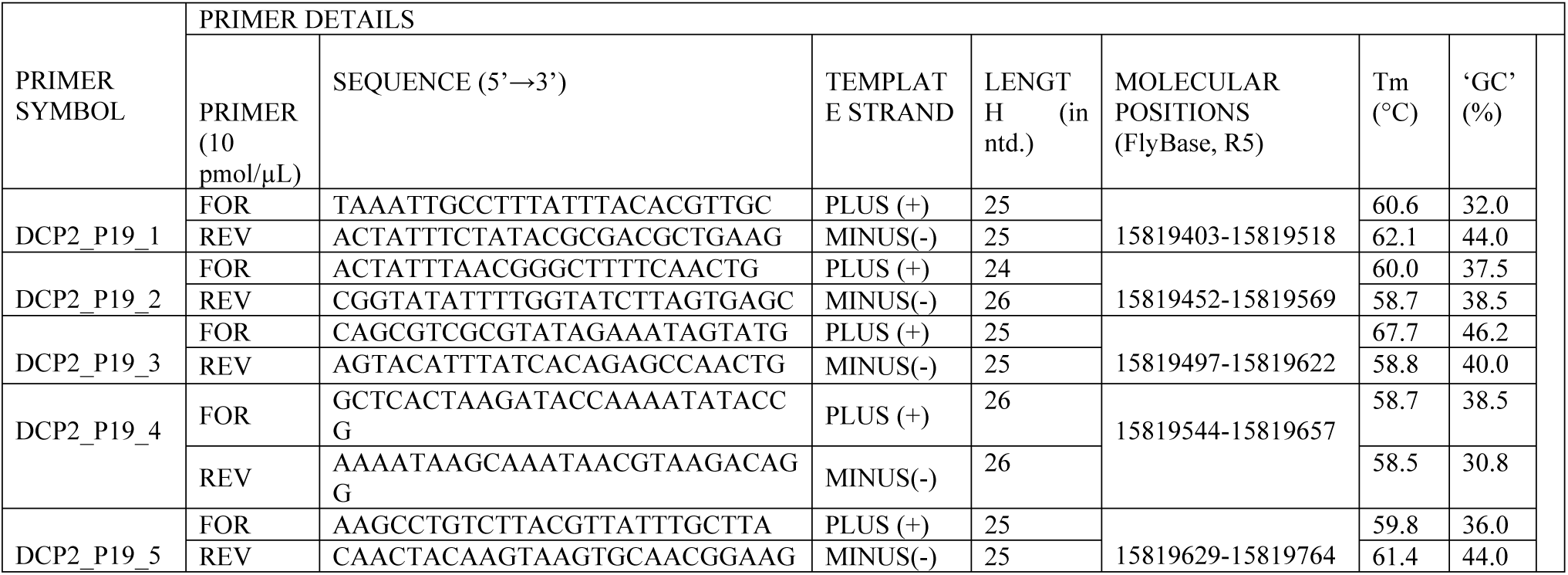
Overlapping set of primers to amplify the genomic region in *DCP2* gene for the region covered by the DCP2_P19 set of primers in the homozygous *l(3)tb* mutant.

**Table S6.**
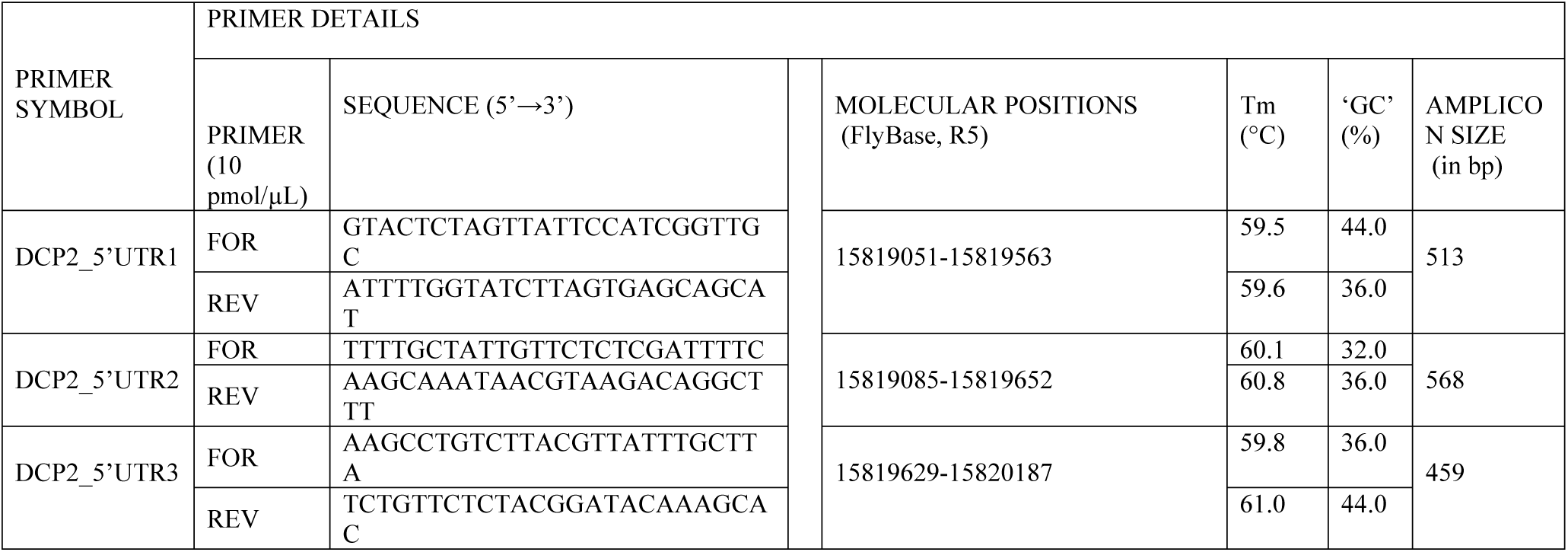
Overlapping set of primers to amplify the complete 5’UTR of genomic region in *DCP2* gene in the homozygous *l(3)tb* mutant. The table also documents the thermal cycler conditions of annealing temperature and extension time for each primer pair. Genomic region amplified by primer pair is also mentioned.

